# Intentionally vs. Spontaneously Prolonged Gaze: A MEG Study of Active Gaze-Based Interaction

**DOI:** 10.1101/2024.12.11.627776

**Authors:** Anatoly N. Vasilyev, Evgeniy P. Svirin, Ignat A. Dubynin, Anna V. Butorina, Yuri O. Nuzhdin, Alexei E. Ossadtchi, Tatiana A. Stroganova, Sergei L. Shishkin

## Abstract

Eye fixations are increasingly employed to control computers through gaze-sensitive interfaces, yet the brain mechanisms supporting this non-visual use of gaze remain poorly understood. In this study, we employed 306-channel magnetoencephalography (MEG) to find out what is specific to brain activity when gaze is used voluntarily for control.

MEG was recorded while participants played a video game controlled by their eye movements. Each move required object selection by fixating it for at least 500 ms. Gaze dwells were classified as intentional if followed by a confirmation gaze on a designated location and as spontaneous otherwise.

We identified both induced oscillatory and sustained phase-locked MEG activity differentiating intentional and spontaneous gaze dwells. Induced power analysis revealed prominent alpha-beta band synchronization (8–30 Hz) localized in the frontal cortex, with location broadly consistent with the frontal eye fields. This synchronization began 500–750 ms before intentional fixation onset and peaked shortly after it, suggesting proactive inhibition of saccadic activity. Sustained evoked responses further distinguished the two conditions, showing gradually rising cortical activation with a maximum at 200 ms post-onset in the inferior temporal cortex during intentional fixations, likely indicative of focused attentional engagement on spatial targets. These findings illuminate the neural dynamics underlying intentional gaze control, shedding light on the roles of proactive inhibitory mechanisms and attentional processes in voluntary behavior.

By leveraging a naturalistic gaze-based interaction paradigm, this study offers a novel framework for investigating voluntary control under free behavior conditions and holds potential applications for enhancing hybrid eye-brain-computer interfaces.

## 1. Introduction

Gaze-based communication and control represent a promising approach for hands-free interaction with computers and other devices, been in development for more than three decades (Duchowski, 2018). It is particularly beneficial for paralyzed individuals (Majaranta, 2011), but also can be used by healthy people, especially as an effective input in virtual, augmented, and mixed reality (VR/AR/MR) (O’Callaghan, 2024; Plopski et al., 2022). To issue a command, a user of this technology typically keeps their gaze on a “gaze-sensitive” element of a graphical interface, such as a screen button, for a relatively short period (Duchowski, 2018; Majaranta, 2011). An eye tracker captures the gaze position, and when the gaze dwell time in the vicinity of a screen button exceeds a specified threshold, the button is activated. The action resembles pressing a computer mouse button or tapping on a touchscreen but involves a temporary suspension of natural spontaneous gaze behavior.

Despite the growing interest in gaze-based human-computer interaction, the brain mechanisms underlying eye movement control in this context have received little attention, likely because this interaction has been considered primarily an engineering challenge. However, gaze-based interaction has unique characteristics that distinguish it from other forms of voluntary behavior. First, the “stillness-based” nature of gaze-based interaction is in stark contrast to typical manual actions, which involve intentional movement. In everyday scenarios, hands remain still by default, whereas the default state for eyes is their frequent motion (primarily through saccades). Secondly, intentional gaze dwells^1^ used in gaze-based interaction represent a rare example of voluntary control executed with minimal effort. Thus, studying the physiological mechanisms specifically related to gaze-based control may enhance this technology. Moreover, using gaze-based actions as a model of voluntary actions could also provide insights into the brain’s mechanisms of voluntary control in general.

Voluntary control of eye movement has been extensively studied in brain research, usually involving salient visual stimuli. In these studies, participants may be required to make voluntary saccades in different directions (often opposite to the stimulus), as in the antisaccade task (Funahashi et al., 1993; Hallett, 1978; Munoz & Everling, 2004), or to maintain a gaze fixation despite an urge to make a saccade, as seen in response inhibition paradigms like the NoGo task (Brown et al., 2006; Brown et al., 2008; Isoda & Hikosaka, 2007) and in delayed antisaccade and prosaccade tasks (e.g., (Ettinger et al., 2008)). These tasks often impose stronger control demands than most everyday activities, which can lead to an overemphasis on the effects of voluntary control and potential confounding with effort-related effects (Jarvstad & Gilchrist, 2019). To address this issue, Jarvstad and Gilchrist (2019) introduced a novel, less demanding paradigm for studying saccade control. Their design included prolonged fixations as a baseline (20 seconds), which still demanded considerable cognitive effort from participants and, therefore, did not fully eliminate unnecessary effort. Moreover, instructions to suppress natural gaze behavior and maintain extended fixation on a target—often to an uncomfortable degree—are common even in baseline conditions of psychological and neurophysiological studies that do not specifically target voluntary eye movement control.

In contrast, gaze-interaction interfaces typically employ much shorter dwell times—often one second or less—to minimize user discomfort. Leveraging gaze-based interaction paradigms could offer valuable insights into the fundamental mechanisms underlying voluntary control, without the need to account for effortful behavior. Voluntary brain control of gaze dwells used for interacting with computers is expected to differ from the control of similarly timed spontaneous dwells, which are highly automatized and basically driven by distinct goals or intentions, or even by reflective reactions. However, since cognitive effort remains minimal even for intentional dwells in this context, any differences in brain activation between intentional and spontaneous dwells during gaze-based interaction are likely to more precisely reflect the voluntary components of eye movement control than traditional paradigms.

Understanding the brain mechanisms underlying low-effort voluntary control of eye movements is important not only for basic neuroscience but also for advancing gaze-based interaction technologies. This knowledge could specifically address the “Midas touch” problem—the challenge faced by gaze-sensitive interfaces in distinguishing intentional eye movement patterns used to issue commands from similar patterns that occur spontaneously or without a clear intent (Jacob, 1990). In particular, when an interface is designed to respond to prolonged dwells, it inevitably risks triggering unintended actions caused by prolonged dwells that are out of conscious control.

Existing countermeasures against the Midas touch problem impose additional effort on users (Majaranta & Bulling, 2014; Velichkovsky et al., 1997). Identifying the differences between intentional and spontaneous gaze dwells could enable the development of brain signal markers capable of distinguishing intentional dwells, allowing interfaces to respond appropriately (Ihme & Zander, 2011; Protzak et al., 2013).

A straightforward approach to studying the difference between intentional and spontaneous gaze dwells is to collect them from participants engaged in tasks using a gaze-based human-computer interface (Shishkin et al., 2016b). The task and visual display should be designed to provoke a sufficient number of spontaneous gaze dwells with a duration exceeding the dwell time threshold used in the interaction. To this end, Shishkin et al. (2016b) developed a gaze-controlled game and recorded electroencephalogram (EEG) data as participants played. Intentional and spontaneous dwells were labeled based on a rule requiring the activation of gaze-based control prior to each move, using a dwell at a designated screen position (the “switch-on button”). Intentional dwells were found to exhibit the EEG phase-locked activity gradually increasing up to the dwell’s termination, likely reflecting anticipation of an interface triggering (Shishkin et al., 2016b). However, this EEG marker did not appear to be specific to the intentionality of a dwell—similar activity could also develop in spontaneous dwells perceived as potentially triggering unintended interface actions, as was likely the case in our online study (Nuzhdin et al., 2017). Unfortunately, the earlier time interval could not be effectively studied due to significant variability in the eye movement patterns prior to the dwell, which was determined using the “switch-on button.”

In the current study, we investigated whether voluntarily prolonged gaze dwells used to control a computer are associated with distinct brain activity that is not present, or at least less pronounced, in spontaneous long dwells from the onset. We employed the same gaze-controlled game as used in the study by (Shishkin et al., 2016b) but modified the gaze-based control rules to minimize the influence of preceding events on the brain activity during intentional dwells. Most of the game rules remained unchanged, as they provided a basis for comfortable interaction and stable motivation in the previous study. Instead of recording the EEG, we co-registered 306-channel magnetoencephalography (MEG) data with eye-tracking data. A special procedure for selecting intentional and spontaneous dwells was developed to ensure contrast on the “voluntariness” scale. The dataset collected in this study has been used previously to assess the usefulness of MEG data for classifying spontaneous and intentional gaze dwells, with a simpler procedure of labeling the intentional and spontaneous dwells (Ovchinnikova et al., 2021).

We hypothesized that certain MEG components would be more pronounced in intentional gaze dwells compared to spontaneous dwells early in their time course, indicating their specific relation to voluntary control. A priori, we could not be certain about the possible locations of such effects; therefore, a whole-head analysis was planned instead of a region-of-interest (ROI)-based one. We identified MEG components phase-locked to dwell onset and non-phase-locked oscillatory MEG components that differed between intentional and spontaneous gaze dwells early in their course.

## 2. Methods

### 2.1. Participants

A total of 32 healthy participants (18-51 years, mean age 25 years; 12 female) were recruited for the study. Among them, seven had previously participated in experiments using the same gaze-controlled game, although with a different sequence of required eye fixations for making moves. The study was conducted in compliance with the Helsinki Declaration and received approval from the ethical committee of Moscow State University of Psychology and Education (protocol 12.03.2015/1). All participants provided written informed consent. During the initial stages of data labeling, three participants (two female and one male) were excluded due to insufficient data (fewer than 50 intentional dwells), leaving a final sample of 29 participants for analysis.

### 2.2. Experimental setup

The experiments were conducted in a magnetically shielded room at the MEG Center (Center for Neurocognitive Research), Moscow State University of Psychology and Education, Russia (http://megmoscow.com/). Participants were seated comfortably in an upright position during the MEG recording. Visual stimuli were presented using a digital light processing projector (Panasonic PT-D7700E-K) with a resolution of 1280 × 1024, projected onto a translucent screen positioned 1.42 meters from the participants’ eyes.

### 2.3. MEG recording

MEG data were recorded using a 306-channel Elekta Neuromag TRIUX™ system (MEGIN/Elekta Oy, Helsinki, Finland) while participants were seated. This system comprises 102 sensor groups, each containing one magnetometer and two planar gradiometers. Before entering the shielded room, four head-position-indicator (HPI) coils were attached to the participant’s head for continuous head position tracking. The positions of the coils and individual anatomical landmarks were digitized using a Polhemus Fastrak system (Polhemus Inc., VT, USA). MEG data were sampled at a rate of 1000 Hz with a frequency range of 0-330 Hz.

To enable synchronization of MEG and eye tracking data, in the beginning and in the end of each game four square signals were sent from the eye tracker server to MEG trigger port. Visual event timing in relation to MEG data was controlled using a photosensor positioned at different locations on the screen at time when no experiments were conducted. The signal from the sensor was recorded along the MEG data and its position relative to the event triggers was checked offline.

### 2.4. Eye tracking and gaze-based control

A MEG-compatible EyeLink 1000 Plus eye tracker (SR Research, Canada) was positioned below the screen, operating in monocular mode at a frame rate of 1000 Hz. It was used both for real-time control of the gaze-controlled game and for recording gaze coordinates for offline analysis. Each 5-minute game session was preceded by a standard 9-point calibration, performed using the built-in SR Research algorithm. Gaze-based control was implemented via the Resonance platform (Nuzhdin, 2019). An online algorithm analyzed gaze coordinates within a 500 ms moving window to detect gaze dwells. A dwell was recorded when two conditions were met:

(a) The range of both X and Y coordinates did not exceed 120 pixels, corresponding to 1.56° of visual angle from the participant’s position (equal to the closest distance between cell centers on the game board), and
(b) No previous dwell had been recorded within 600 ms of the new dwell.

These prolonged dwells functioned similarly to mouse clicks or touchscreen taps in the traditional Lines game, triggering specific events based on various game conditions (see game description below).

### 2.5. The eye-controlled game

To simulate the natural behavior of users interacting with a gaze-based system, the experiment had participants freely play a specially designed eye-controlled game, EyeLines, originally described by (Shishkin et al., 2016b) and modified for the purposes of this study. The game was designed to elicit two distinct types of gaze behavior, both involving dwells (500 ms or longer): intentional gaze use for making moves, and spontaneous oculomotor activity (including visual exploration of the game board and deliberation on future moves without active interaction). This setup enabled the collection of a sufficient number of both intentional and spontaneous dwells. The game rules and software were structured to ensure that a significant proportion of long dwells could be reliably classified as either intentional or spontaneous. The classification of dwells into these two categories, along with the comparison of corresponding gaze and MEG data, was conducted during offline data analysis.

The EyeLines game board, displayed on a white background rectangular area (413 × 330 mm, 16.6° × 13.3°), was projected onto the screen (Figure 1). As in (Shishkin et al., 2016b), the board consisted of 7×7 square grey cells with equal gaps between them (in contrast to the original Lines game, typically played on a 9×9 board with a slightly different visual layout; these changes were introduced by Shishkin et al. (2016) to enhance gaze-based control). The distance between the centers of adjacent cells was 1.6°. An additional single cell, used to confirm gaze-based selections, was placed to the left of the board in half of the games and to the right (as shown in Figure 1) in the other half of the games.

**Figure 1.**
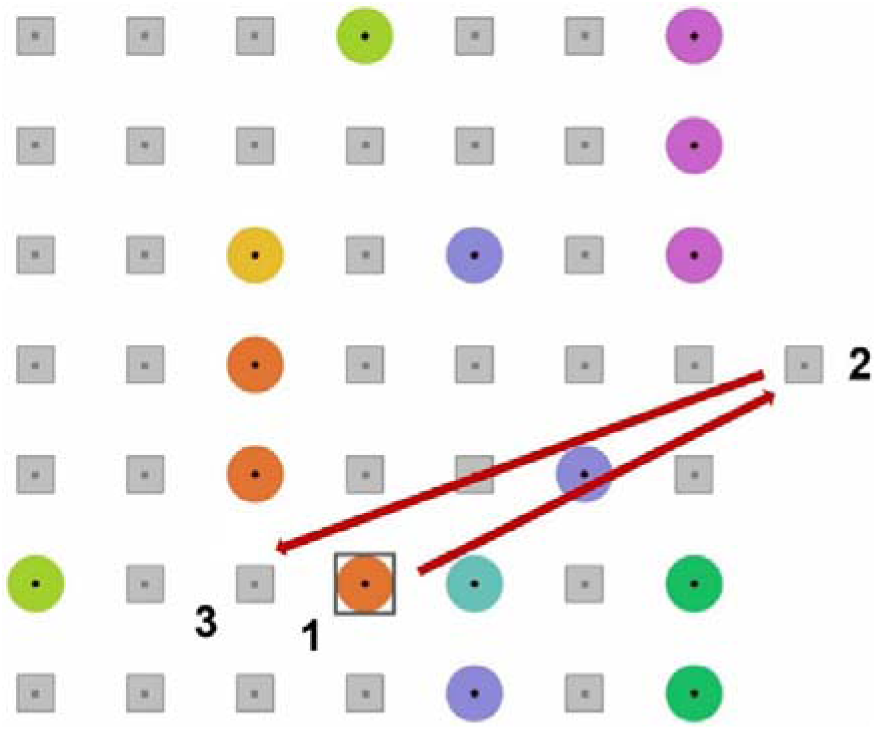
An example of *EyeLines* game position (Ovchinnikova et al., 2021, license CC BY). Here, a screenshot of an actual position is presented, with digits and arrows added to explain the sequence of gaze dwells required to make a move: 1 – dwell for selecting a “ball” (after 500 ms dwell, a square frame appeared around the ball), 2 – dwell on a remote “button” confirming the selection (after 500 ms dwell, a ball appeared over this cell), 3 – dwell on a free cell (after 500 ms dwell, the ball in cell 1 disappears and appears in cell 3). The distance between cell centers was 1.6°.

A participant’s objective in the EyeLines game was to score as many points as possible. Points were awarded each time a participant successfully aligned four or more “balls” (circles) of the same color in a straight line. When such a line was formed, all balls within it disappeared from the board. After any other move, as well as at the start of each game, three colored balls appeared in randomly selected free cells on the board. Consequently, if a participant did not frequently create lines of four or more balls, the board would quickly fill up, leaving fewer available cells for new balls. The game ended either when fewer than three free cells remained, leaving no space for new balls, or after five minutes had passed without reaching this condition. At the end of the game, the participant’s score was displayed.

The color of each new ball on the board was randomly chosen from seven possible colors. A different color was used for the ball that appeared in the confirmation button position. To aid in fixation accuracy, black dots were placed at the centers of balls and empty cells.

As in the original Lines game, each move in EyeLines involved selecting a ball and then selecting a free cell to which it should be moved. Moves were only possible if a “free path” of adjacent free cells existed between the ball’s initial and new positions. Unlike the original Lines game, in EyeLines, selections of both the ball and the new cell were made using gaze dwells (see above the Eye tracking and gaze-based control section for details). Ball selections had to be confirmed by fixating on a confirmatory “button,” a cell located outside the board (labeled as position 2 in Figure 1; hereafter referred to as “the button,” as this term was used to explain the move process to participants).

Participants were instructed to execute each move by following this sequence of actions:

(a) Identify the ball they wish to move and the free cell to which they intend to move it.
(b) Fixate on the ball until a square frame appears around it (“ball pre-selection,” labeled as position 1 in Figure 1).
(c) Fixate on the button (position 2 in Figure 1) until a ball of a different color appears within it, indicating that the selection has been confirmed (“selection confirmation”).
(d) Fixate on the free cell where the ball is to be moved (position 3 in Figure 1) until the ball is “moved” to it (“ball movement”).

The “ball movement” process involved several simultaneous events: the selected ball and the frame around it disappeared from position 1, the ball indicating the selection disappeared from position 2, and the “moved” ball appeared in position 3, maintaining the same color as the ball originally in position 1.

All events—“ball pre-selection,” “selection confirmation,” and those comprising “ball movement”— were triggered by 500 ms dwells detected online, as explained in the previous subsection, under the additional condition that the median gaze coordinates within the 500 ms time window fell within a 120 x 120 pixel area (1.6 x 1.6°), centered on the related ball, button, or free cell. (Instead of two dwell time thresholds, 500 ms and 1000 ms, used by (Shishkin et al., 2016b), we only used only the shortest one, as it allowed for the collection of a sufficient number of intentional and spontaneous gaze dwells of similar duration).

Participants were instructed that if a frame appeared unexpectedly around a ball they did not intend to select, they should simply ignore it, as only the most recently pre-selected ball could be confirmed via a button dwell. They could change their pre-selected ball by dwelling for 500 ms on any other ball. In such cases, the frame would disappear from around the previously pre-selected ball and reappear around the newly pre-selected ball. There were no limitations on the number of selection changes.

In data analysis (described below), ball pre-selections followed by a dwell on another ball rather than a button dwell were categorized as spontaneous dwell. In contrast, quickly confirmed ball pre-selections were classified as intentional dwells. These details were not disclosed to participants; they were only informed that the purpose of the experiment was to study brain activity during gaze-based interaction with a computer.

The requirement of pre-selection confirmation via a button dwell differed from the game design used in (Shishkin et al., 2016b). In that study, a button dwell activated control, allowing immediate selection of a ball through subsequent dwells. Spontaneous dwells were defined as those occurring without prior activation of control. However, in that design, a high-amplitude saccade from the activation button preceded intentional dwells, introducing “contamination” from saccade-related brain and oculographic activity. Additionally, the amplitude of the preceding saccade strongly influences brain activity early in the course of fixation (e.g., (Nikolaev et al., 2016)). Therefore, the game design was modified to equalize, as much as possible, the onset conditions for intentionally and spontaneously prolonged fixations, minimizing the influence of factors unrelated to voluntary control.

### 2.6. Procedure

Participants received instructions and engaged in at least two practice games with the confirmation button positioned on both the left and right sides. Additional adjustments of the eye tracker, further instructions, and/or additional practice sessions were provided if necessary. Once the participant demonstrated adequate gaze-based control and a clear understanding of the game rules, the main series of eight games commenced. Eye tracker calibration was performed before each game, including the practice sessions. Participants were allowed a rest period of 1-3 minutes between games. The experimenter continuously monitored the participant’s gaze, which was displayed as a dot moving over a copy of the game board. If signs of poor control were observed (an infrequent occurrence, happening no more than once per participant), the game was halted, its data discarded, and a new calibration was conducted before restarting the game.

All eight games were played under identical conditions for all participants, with the only variable being the position of the confirmation button. The button’s position for the first game was randomized and counterbalanced across participants. In subsequent games, the button position alternated between symmetrical left and right positions (Figure 1 provides an example of the right button position).

### 2.7. Gaze data analysis

The primary focus of the analysis was on gaze dwells lasting longer than 500 ms on balls, which led to ball pre-selection during the game. Preprocessing of online-detected dwells involved the following steps:

(a) Rejection: Dwells longer than 500 ms were rejected if gaze coordinates were partially lost (e.g., due to blinks detected online by the SR Research algorithm, or eye tracker malfunctions) or if the participant violated instructions or game rules (e.g., dwell on a free cell without prior selection confirmation, or attempting to move a ball to a cell with no available path).
(b) Onset correction: The onset time of the dwell and the time of leaving the fixated object were corrected. This step involved identifying an eye fixation detected by the built-in eye tracker algorithm, which matched the > 500 ms dwell identified by the EyeLines algorithm (using criteria for saccades: acceleration > 8000°/s², speed > 30°/s, and a minimum duration of 6 ms). If no fixation was detected by the eye tracker within 50 ms of the EyeLines dwell onset, the latter was discarded (occurring in less than 5% of cases). Otherwise, the time of the closest fixation identified by the eye tracker was used as the corrected dwell onset. The dispersion-based algorithm was preferred for online interaction due to its perceived naturalness and effectiveness in pilot studies, while the built-in eye tracker criterion provided the precise moment of gaze stop, which was crucial for MEG data analysis. This procedure effectively excluded most dwells with incorrectly defined onsets due to slow response times in the online interaction algorithm (e.g., calibration issues).
(c) Labelling: The identification of spontaneous and intentional dwells was primarily determined by subsequent events, such as dwell on another ball or the confirmation button. However, additional filtering of dwells associated with specific gaze behavior patterns was necessary to ensure a high level of certainty regarding strictly intentional dwells (i.e., those intended from the outset). The selection of dwells for analysis was based on the alignment between the eye movement pattern and the expected behavior of the player during spontaneous or intentional ball selection. Specifically, an intentional ball selection should be promptly confirmed by activating the button, ultimately leading to the execution of a move. To evaluate this, we analyzed the delay in the participant’s response to feedback from the selected ball (the time from ball selection to the initiation of a saccade) and the time taken to shift gaze to the confirmation button (the time from the first saccade after selecting the ball to the onset of the dwell on the button). For most participants, the mode of the reaction time histogram fell within the range of 270-320 ms, while for others, it was shifted to approximately 400 ms explained by the more frequent occurrence of delayed reactions. To capture the most representative reaction time interval, a threshold of 500 ms was established for reaction speed to the ball’s feedback to identify intentional dwells. For the time required to shift gaze to the button, a threshold of 200 ms was chosen, which exceeds the typical duration of a single saccade but is insufficient to allow for additional dwells that could indicate a search for a favorable spot for a spontaneously selected ball. The main criterion for identifying spontaneous selections was the presence of at least one unconfirmed selection of another ball. This ensured that the chosen dwell was not a result of calibration or activation issues, which could have led to a missed selection of the intended ball.

### 2.8. MEG preprocessing

Sensors with contaminated signals were manually excluded, and the MaxFilter routine (MEGIN/Elekta Oy, Helsinki, Finland) was applied using the following settings: temporal signal space separation (tSSS) (Taulu & Simola, 2006) with t = 10 s and corr = 0.9, movement correction (MaxMove), and head origin normalization to the standard position (for details, see MaxFilter User’s Guide, https://ohba-analysis.github.io/osl-docs/downloads/maxfilter_user_guide.pdf).

Synchronization of MEG and eye tracker data was achieved for each game by adjusting the MEG time so that the synchronization triggers in both datasets coincided. MEG recordings were segmented into epochs of [-1000 2000] ms relative to the onset of spontaneous and intentional dwells. MEG analysis was divided into two pipelines aimed at investigating evoked power (phase-locked event-related fields) and induced power (non-phase-locked activity in the alpha and beta bands).

### 2.9. Induced power analysis

To explore the oscillatory dynamics during both spontaneous and intentional dwells, we employed the method of simultaneous diagonalization of covariance matrices of the signal in compared conditions (generalized eigendecomposion, GED – see also (Cohen, 2022; Nordhausen & Ruiz-Gazen, 2022; Parra & Sajda, 2003)). This method enables the analysis of multidimensional data (in our case – MEG) by utilizing hypothesis-driven spatial filters to enhance distinctions in the power of signal frequency bands, while concurrently minimizing irrelevant activity and reducing data dimensionality. Our hypothesis posited distinct manifestations between intentional and spontaneous dwells, specifically focusing on variations in levels of disinhibition within the eye motor system (as indicated by central alpha(mu) and beta rhythms), and the divergent engagement of the visual analyzer (potentially, leading to alterations in occipital alpha). We anticipated intentional dwells to demonstrate controlled inhibition in eye movement, whereas spontaneous dwells were expected to engage in simultaneous visual information processing and saccadic activity.

Therefore, our investigation focused on combined alpha+beta frequency range (8–30 Hz) within the interval [-0.25 – 0.4] seconds relative to fixation onset, which encompassed the saccade and the primary phase of the dwell.

While individual analysis is typically more sensitive, it presents challenges for interpretation at the group level due to difficulties when matching corresponding components across individual participants. Therefore, for this analysis, we opted for a less sensitive method involving pooling signals from all participants and the computation of group-level spatial filters (components). In accordance with the recommendations outlined in (Cohen, 2022), the following algorithm was employed for analysis:

1) We equalized the number of participant’s spontaneous and intentional epochs by randomly selecting a maximum of 100 epochs per condition and per participant. If a condition had fewer than 100 epochs in a perticipant, we equalized the number of epochs per condition by randomly removing epochs from another condition of this participant. As a result of this procedure, 20 participants had between 95-100 epochs per condition, while the remaining participants had between 50-80 epochs per condition, resulting in a total of 2560 epochs per condition for the whole group. No participant had less than 50 epochs per condition.
2) To reduce contribution of evoked components, MEG signals were averaged over epochs and subtracted from each epoch. This procedure was done separately for spontaneous and intentional epochs and for each participant. Epoch signals were normalized for each participant using the median absolute deviation (MAD), ensuring that across all gradiometers and epochs within the [-2 to 2] s window relative to fixation onset, the MAD was 1. The same normalization was applied separately to all magnetometers.
3) Signals were bandpass-filtered in 8-30 Hz frequency range using FIR zero-phase filter. Following that, dimensionality reduction of the data was performed using PCA retaining only components that explained at least 0.001% of the variance (resulting in a total of 77 components), ensuring that the data were full rank.
4) Covariance matrices were computed: one for each condition for all participants collectively (subsequently denoted as S for intentional trials and R for spontaneous trials) followed by generalized eigendecomposition (GED) computation expressed by the following formula:

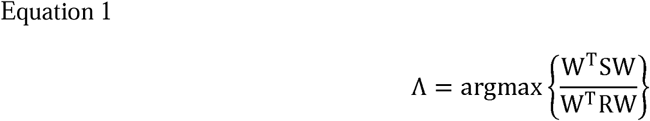

Where Λ denotes the eigenvalues of the SR^-1^ matrix, the columns of W represent the spatial filters (inverse operators) for de-/synchronization components, and W^-T^ represents forward models of the components (Haufe et al., 2014).

5) To mitigate overfitting, step 4) was repeated 10,000 times with trials’ label permutation. We retained columns of W whose eigenvalues exceeding the 1st percentile (for the first components) or the 99th percentile (for the last components) of the permutation distribution based on comparisons between the original Λ obtained and the distribution obtained through permutations.
6) For epochs projected onto the selected components, time-frequency wavelet transform was performed using the Morlet wavelet with a variable number of cycles following the procedure proposed by (Moca et al., 2021). Wavelet scales were linearly spaced from 3 to 45 Hz with 0.5 Hz step, minimum number of cycles was set to 5 with 1 to 30 superresolution order as described by Moca et al., power values were converted to decibel. Spectrograms were generated by averaging all trials with trimmed mean with 10% of marginal values removed, followed by either calculating difference between conditions, or baseline subtraction represented by mean value within [-1.5 –-1] s interval prior to fixation onset.
7) To evaluate the statistical significance of differences in synchronization levels among oscillatory components, we utilized differential spectrograms contrasting intentional and spontaneous conditions. Statistical significance was determined through a permutation cluster test. Namely, the differential spectrograms were computed for randomized partitioning into intentional and spontaneous trials, repeated 10,000 times; all differential spectrograms were converted into binary matrices, with a value of 1 assigned to data falling within the marginal 1% distributions derived from permutations and 0 for the rest of the data; a threshold for the size of the largest cluster (contiguous area of «1» pixels) was determined as the 99^th^ percentile of encountered on permuted maps. Finally, these thresholds were applied to the differential spectrogram maps based on true label partitioning, defining statistically significant time-frequency clusters.

This procedure was also separately applied to alpha (8–15 Hz) and beta (15–30 Hz) ranges as well as pre-and post-onset time intervals ([-0.4 – 0], [0 – 0.4] seconds). In these cases, only shifts in statistical significance of spatial components and band power statistics were obtained, while the whole pattern of the results followed what was revealed in the broad band analysis. To save space, we do not report these results.

### 2.10. Evoked fields analysis

The multivariate approach of extracting significant spatial projections, which we employed in the analysis of induced activity, can also be used in the analysis of phase-locked, evoked signal components (Cohen, 2022; de Cheveigné & Parra, 2014). Two adaptations for evoked activity were proposed, denoising source separation (DSS) (Särelä et al., 2005)) and Fischer-criterion beamforming (FCB) (Pires et al., 2011). They were applied for multidimensional EEG and MEG analysis in a number of studies (Das et al., 2020; Molloy et al., 2019; Presacco et al., 2016). These methods are effectively a dimensionality reduction procedure with output components optimized according to two criteria: reducing within-condition variance (i.e., isolating phased-locked responses and suppressing non-phased-locked activity) while simultaneously increasing between-condition variance (i.e., differences between compared conditions). A small distinction between DSS and FCB lies in the fact that in DSS, the optimization steps for evoked activity and differences between conditions are separated, while FCB addresses them simultaneously.

We applied FCB to test the hypothesis that evoked activity differs between compared conditions: spontaneous and intentional gaze delays. Similar to the algorithm utilized for analyzing induced activity, the following algorithm was employed for evoked fields:

1) The same epochs were selected (2560 per condition) as for induced activity analysis (see step (1) in induced analysis).
2) Epochs underwent a 20 Hz low-pass filtering using FIR zero-phase filter, baseline correction to the interval [-0.3 –-0.1] s, and normalization to the overall standard deviation of the same baseline interval across all epochs within the same participant.
3) The Fischer-criterion beamformer was computed for the interval [0 – 0.4] s. Unlike for induced activity, the pre-fixation period was not considered to avoid signal contamination from saccadic artifacts. The Fischer criterion is solved as follows (Pires et al., 2011):

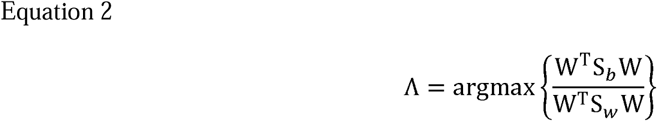

Where Λ denotes the eigenvalues of the *S_b_S_w_*^-1^ matrix, *S_b_* represents the between-condition matrix, and *S_W_* represents the within-condition matrix, computed from

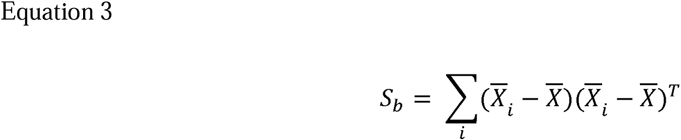

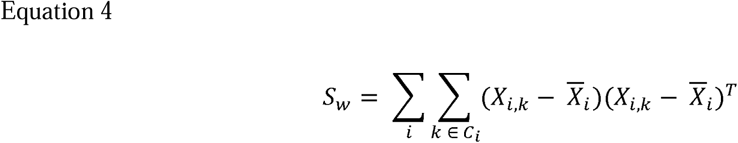

where 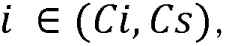 with Ci denoting the intentional condition, Cs denoting the spontaneous condition, and 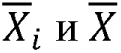 representing the arithmetic average across a condition or across all epochs.

4) Similar to induced field analysis, step 3) was repeated 10,000 times with trials’ label permutation, retaining only components with Λ above the 99^th^ percentile of the permutation distribution.
5) Epochs were projected onto the selected components and rescaled to the standard deviation of the baseline interval [-0.3 –-0.1] s as in step 2), and a 99% confidence interval was evaluated for differential ERF using permutation test (Efron & Tibshirani, 1994). To achieve this, differential evoked fields, after using median averaging per condition and participant, were obtained with condition label permutation (10,000 times), their amplitude was thresholded with a 1% threshold from the distribution, and continuous segments (time-clusters) above the threshold were filtered based on a 1% magnitude from the clusters observed for the recombinant distribution.
6) Selected components from step 4) were projected back onto the sensor space for source-level visualization, enabling a collective analysis irrespective of their amplitude while considering their mixing.

### 2.11. Source-level analysis

Due to unavailability of participants’ MRI data, Colin27 brain template (Holmes et al., 1998) was used to compute cortical activation patterns. The cortical surface meshes comprised 10,000 vertices (sources) across both hemispheres. Statistical inference in this study was conducted on sensor-level data, however, employing source localization techniques could yield valuable insights into the specific brain regions underlying the observed evoked and induced dynamics.

Forward model computation was performed with Brainstorm (Tadel et al., 2011), which is documented and freely available for download online under the GNU general public license (http://neuroimage.usc.edu/brainstorm). The forward solution was calculated using overlapping spheres approach (Huang et al., 1999). The inverse solution was computed by Brainstorm built-in minimum norm estimation algorithm applied with the default settings (“kernel only” as the output mode, three as the signal-to-noise ratio, the source orientation constrained to perpendicular to the cortical surface, the depth weighting restricting source locations to the cortical surface and the whitening PCA). A noise covariance matrix, necessary to control noise effects on the solution (Bouhamidi & Jbilou, 2007), was calculated either from empty room recording (induced analysis) or from baseline interval for evoked analysis.

For both evoked and induced analyses, the group-level pattern (filter’s forward model) of each significant component was multiplied by individual inverse operators, normalized for each participant (ensuring the sum of absolute values across all sources was set to 1), and subsequently averaged across participants in the source space. Sources located on the medial cortical surface, including the corpus callosum and cingulate cortex, were excluded. Retained vertices—those with weights exceeding 50% of the maximum source magnitude (corresponding to the top 6.5% for induced components and 9–11% for evoked components)—were clustered to retain only spatially distinct activations, defined as clusters with more than 100 sources and inter-source distances of less than 1 cm. These activations were visualized on cortical surface meshes. To determine the anatomical locations of the components, the weighted centroids of source clusters were calculated in MNI-normalized space. To ensure accuracy, clusters were constrained to remain within a single hemisphere and limited to either the dorsal or ventral surface of the cortex. This approach prevented centroids from being positioned deep within or between hemispheres.

### 2.12. Control for eye movement artifact contribution

Eye movements can significantly differ between spontaneous and intentional dwells, and related artifacts are common in brain signals co-registered with eye tracker data (Nikolaev et al., 2016). In our evoked MEG data analysis, we employed a signal separation procedure that itself provided some level rejection of eye movement-related interference. However, synchronizing MEG data with eye movements may still increase the likelihood of eye movement artifacts contaminating the MEG signal, making it essential to verify the potential contribution of such artifacts to the observed MEG components. We selected trials preceded by a shift in either the OX or OY coordinates of plus or minus 200 pixels (i.e. right, left, up and down) within a 100 ms window around dwell onset (i.e., saccades with amplitudes larger than 2.6°). We then averaged their evoked fields and eye tracker data, visually inspecting the resulting plots for any signs of eye movement artifacts contributing to the evoked fields. For analysis of induced activity, spectrograms were examined for the presence of low-frequency-dominated broadband activity, characteristic of blink and saccade artifacts. A secondary check, applicable to both induced and evoked components, involved inspecting the patterns (forward models of separating filters) for any contribution to the signal from the frontal sensors.

## 3. Results

### 3.1. Data labelling

On average, the game consisted of 36±7.6 moves (mean±SD), with an average time between moves of 8.28±1.78 seconds. Ball selections (including unconfirmed ones) occurred every 3.35±0.50 seconds. Detailed information for each participant can be found in Table S1. The labeling of dwells that led to ball selections, according to the criteria described in the Methods (see section Gaze Data Analysis), revealed a 2:5 imbalance in favor of spontaneous dwells. The total number of dwells that met all the eye-tracking data quality criteria was 12,023 across 29 participants. In the next step, dwells with abnormal MEG signals were excluded, and the number of spontaneous and intentional dwells was balanced for each participant, leaving a total of 6,748 dwells with a 1:1 ratio between spontaneous and intentional. For the MEG analysis, the number of dwells was further reduced to ensure equal contribution from each participant to the group analysis, resulting in a total of 5,120 dwells. For all subsets, the dwells corresponding to games with the button positioned on the right and left were balanced, with a 52:48 split for the whole set. The number of dwells per participant in the raw data, at different stages of data pruning and in the final dataset is presented in Figure S1.

### 3.2. Gaze behavior

Labeling dwells as spontaneous and intentional was based on different gaze behavior patterns prior to and after the dwells (roughly illustrated in Figure 2A). Therefore, it was not unexpected that certain eye movement characteristics differed dramatically between intentional and spontaneous dwells. However, the distance of the gaze from the center of the dwell (median over the interval 0-500 ms; the left panel of Figure 2B) had almost identical time courses for intentional and spontaneous gaze dwells in the first 200-250 ms. The corrective saccade typically occurred at the same time (around 200 ms) and had similar amplitude in the two types of dwells. After 250 ms, in the case of intentional dwells, gaze movements were restricted to a small area (0.1°) around the center. In contrast, for spontaneous dwells, the gaze rapidly shifted away from the center; as we analyzed only cases where the gaze remained within the “sensitive” area of the ball until 500 ms, up to this time the deviation was moderate “by definition”. Change in distance to the ball’s center (the right panel of Figure 2B) indicated that during intentional dwells, the gaze progressively moved closer to the center, whereas during spontaneous dwells, the gaze drifted away, likely due to attention shifting to surrounding elements of interest. Additionally, intentional dwells were characterized by a synchronous saccade around 750 ms, directed toward the confirmation button. This 750 ms point, i.e., about 250 ms after the onset of the visual feedback indicating ball selection, should correspond to a time when participants were able to react to the feedback. As a result, dwell time (duration) distribution beyond the 500 ms threshold highly differed between intentional and spontaneous dwell (Figure 2C, left).

**Figure 2.**
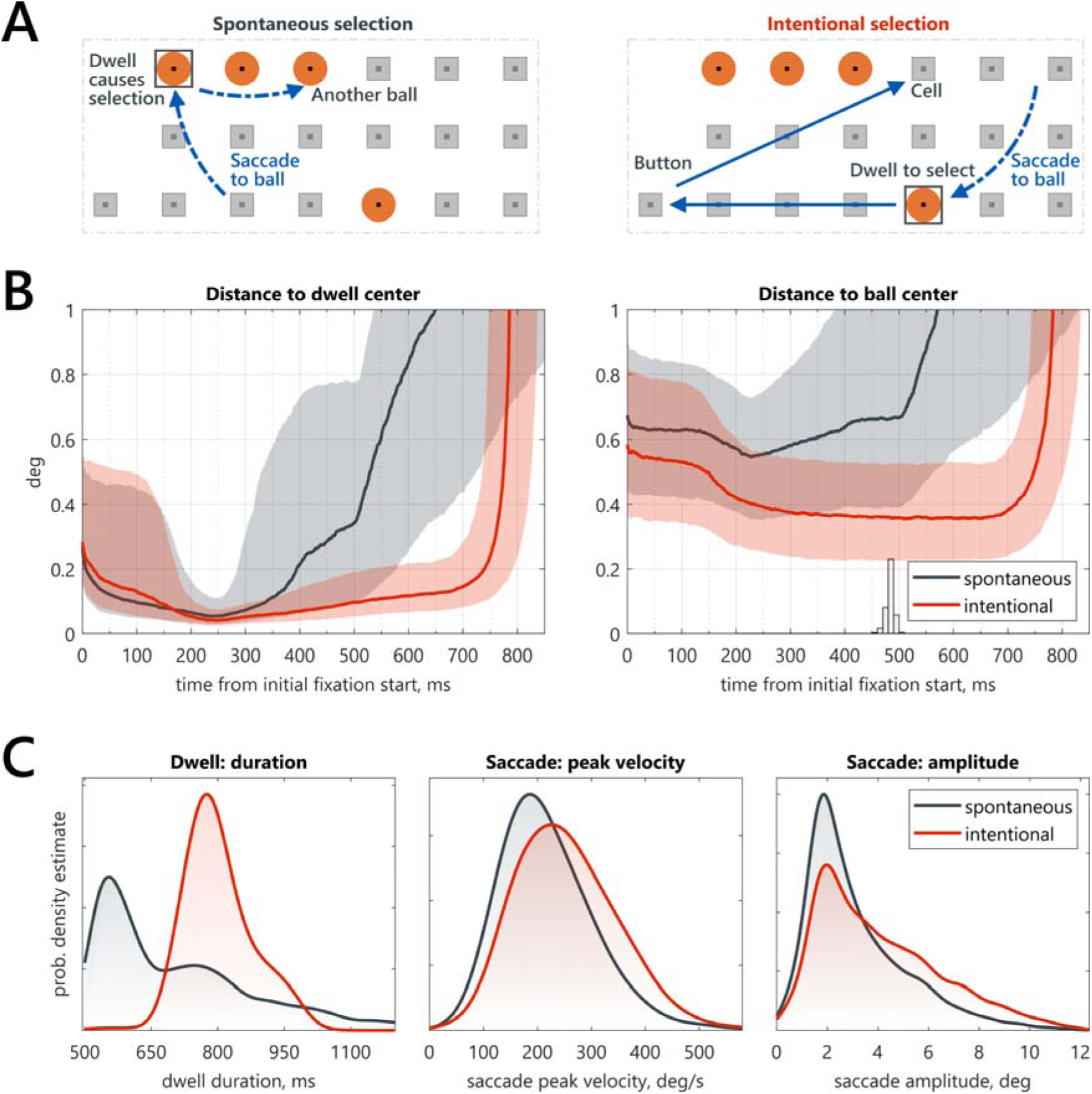
Eye movement activity during spontaneous and intentional dwells. A – Example of dwell and saccade sequence during spontaneous (left) and intentional (right) ball selections. B – Median time courses (solid lines) and interquartile ranges (IQR, shaded area) for gaze deviation from the center of the dwell (left) and from the center of the target ball (right). Black indicates spontaneous dwells, and red indicates intentional dwells (n = 3374 per category). Feedback onset time distribution is shown as a histogram in the right panel. C – Histograms showing the distributions of the following metrics (from left to right): total dwell duration, peak velocity and amplitude of the dwell-preceding saccade towards the ball. Black indicates spontaneous dwells, and red indicates intentional dwells (n = 3374 per category).

When examining the characteristics of the saccades preceding the dwell, only slight and statistically insignificant differences were observed between the conditions. There was a trend toward higher peak velocity (p = 0.071; Figure 2C, center) and amplitude (p = 0.078; Figure 2C, right) in the intentional condition, likely due to some pre-planned selections of distant balls.

### 3.3. MEG: Induced power

The permutation test applied to GED components revealed two significant spatial components (p = 0.0034 and p = 0.0053), which differentiated intentional from spontaneous dwells (their sensor-level topographies are shown in Figure 3, top panel). These components exhibited pre-central scalp topography, lacked left-right asymmetry, and were marked by pronounced event-related synchronization (ERS) within the alpha and beta ranges of MEG (Figure 3, bottom panel). This synchronization began approximately 0.75 seconds before the onset of fixation, intensified around the onset, and peaked approximately 0.25 seconds afterward.

**Figure 3.**
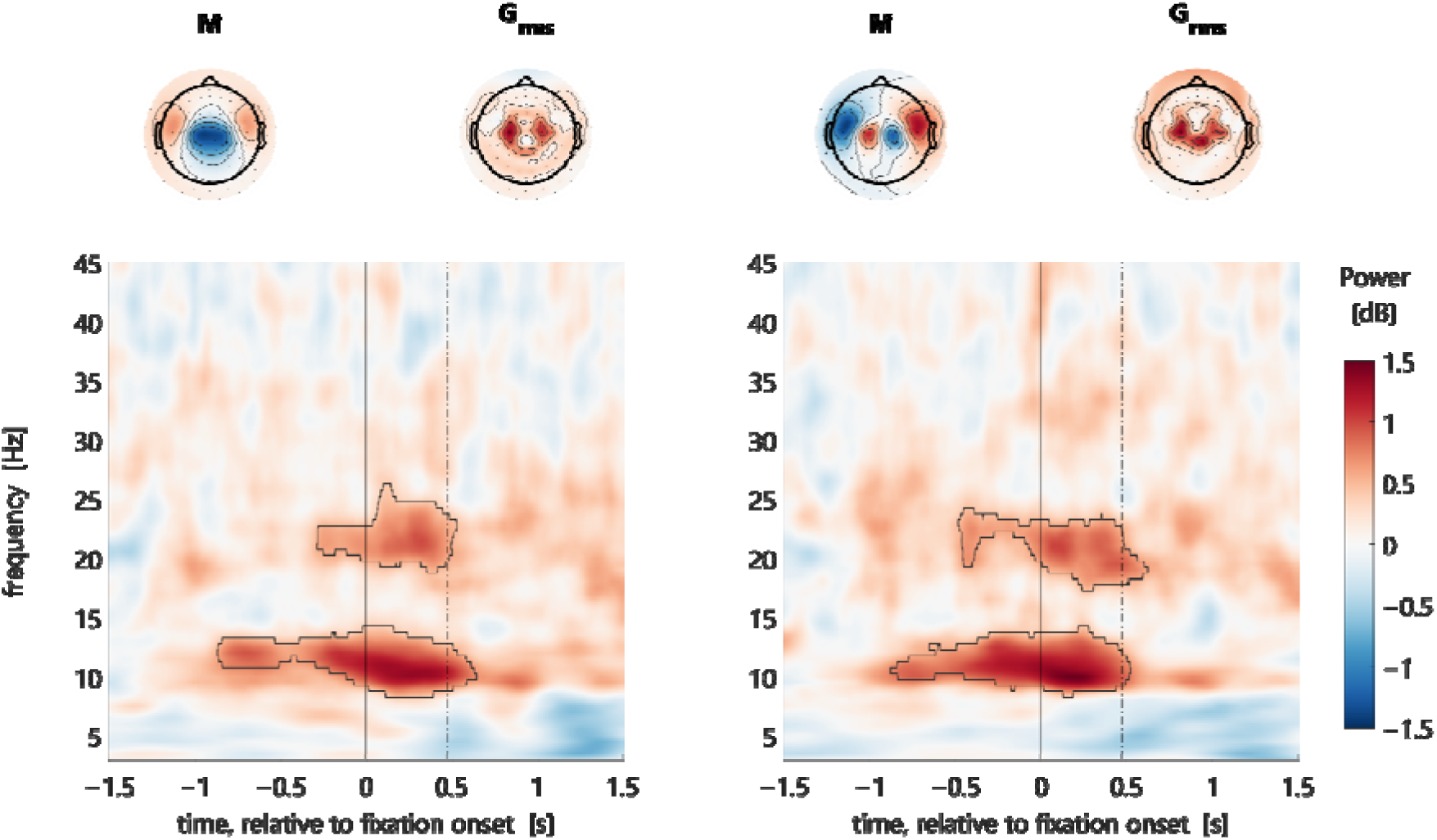
Distinct Oscillatory Signatures of Intentional vs. Spontaneous Gaze Dwells. Left and right panels depict the two distinct induced field components differentiating intentional from spontaneous dwells. At each panel, the top row displays the components projected onto magnetometers (M) and the root mean square of planar gradiometers (G_rms_). The bottom row shows the respective time-frequency maps for each component. Statistically significant clusters outlined by solid black lines (p < 0.01). Vertical lines at 0 and 0.5 s show dwell onset and approximate feedback time, respectively.

Based on the dipole structure inferred from the components’ forward models, it appeared that each component was generated by two closely positioned dipoles with different orientations (Figure 3, upper panel). Time-frequency maps of the components revealed that differential alpha and beta synchronization did not persist throughout the entire dwell duration (∼750 ms) but nearly disappeared around 500 ms after dwell onset (Figure 3, bottom panel).

Examining the time-frequency dynamics of the differential components separately for spontaneous and intentional dwells (Figure S2) suggests that the observed differences in oscillatory activity may stem from two distinct processes—enhanced baseline-normalized synchronization or markedly reduced desynchronization in intentional compared to spontaneous dwells. However, in our free-behavior paradigm, accurate baselined normalization poses a challenge due to the rapid succession of intentional and spontaneous gaze dwells. Due to this limitation, our statistical analysis focused solely on the differential spectrograms, which formed the basis for our conclusions regarding the impact of intentional dwells on oscillatory activity.

Source-level analysis revealed that the cortical localization of two primary components has a substantial overlap. The components were located within the motor, premotor, and superior frontal cortices, including regions such as the frontal eye field (FEF) and supplementary eye field (SEF) (see Figure 4 and Table S2).

**Figure 4.**
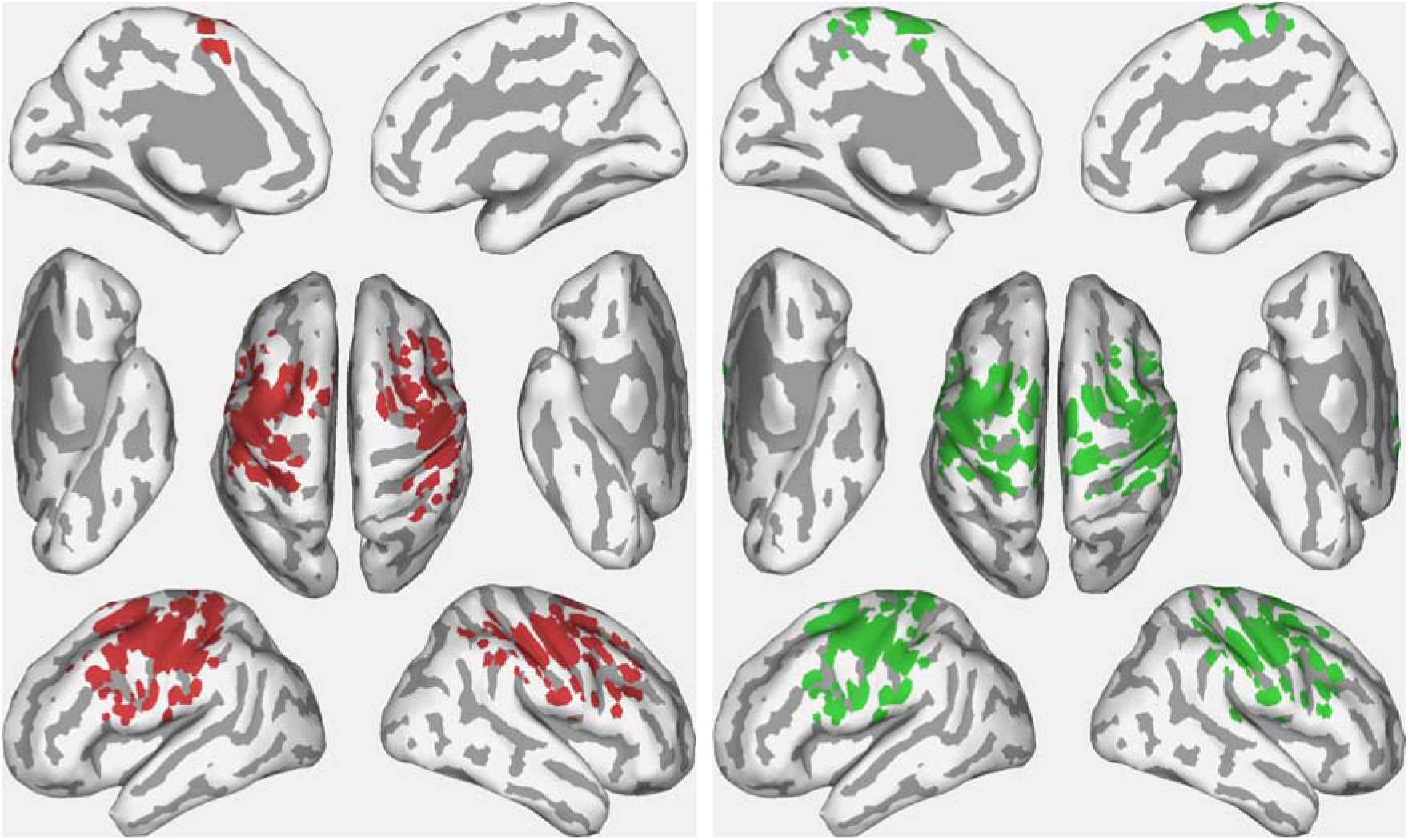
Source representation of group-level spatial patterns for oscillatory activity. Group-averaged source representations of two significant group-level spatial patterns (forward models of the components, see Fig. 3) for oscillatory activity in the alpha-beta frequency range (8–30 Hz). Inverse solutions were constructed using the Colin27 brain template and individual noise covariance. Only the clustered cortical sources with values above half-maximum across the entire cortex are shown (see Methods for details): red for the first spatial component and green for the second. Dark areas correspond to cortical sulci, while light gray areas represent gyri.

### 3.4. MEG: Evoked fields

Analysis using the Fisher criterion beamformer and subsequent eigenvalue significance testing identified three significant components differentiating intentional from spontaneous gaze dwells (label permutation tests yielded p < 0.0001, p < 0.0001, p = 0.0096, respectively) (Figure 5). The first spatial component, corresponding to the largest eigenvalue—which reflects the greatest contribution to differential evoked activity—demonstrated a sustained deflection in the evoked magnetic field under the intentional condition. This deflection began approximately 100 ms before fixation onset, peaked around 200 ms post-onset, and persisted throughout the dwell and until the subsequent intentional action (button activation). The second component displayed an initial transient positive deflection from ∼0.05 to 0.08 s post-onset, followed by a prolonged deflection of opposite polarity from 0.35 to 0.75 s. The third component was transient, primarily tied to fixation onset, and coincided with both the initiation of the intentional dwell and the subsequent intentional confirmatory saccade (from 0.7 to 0.8 s).

**Figure 5.**
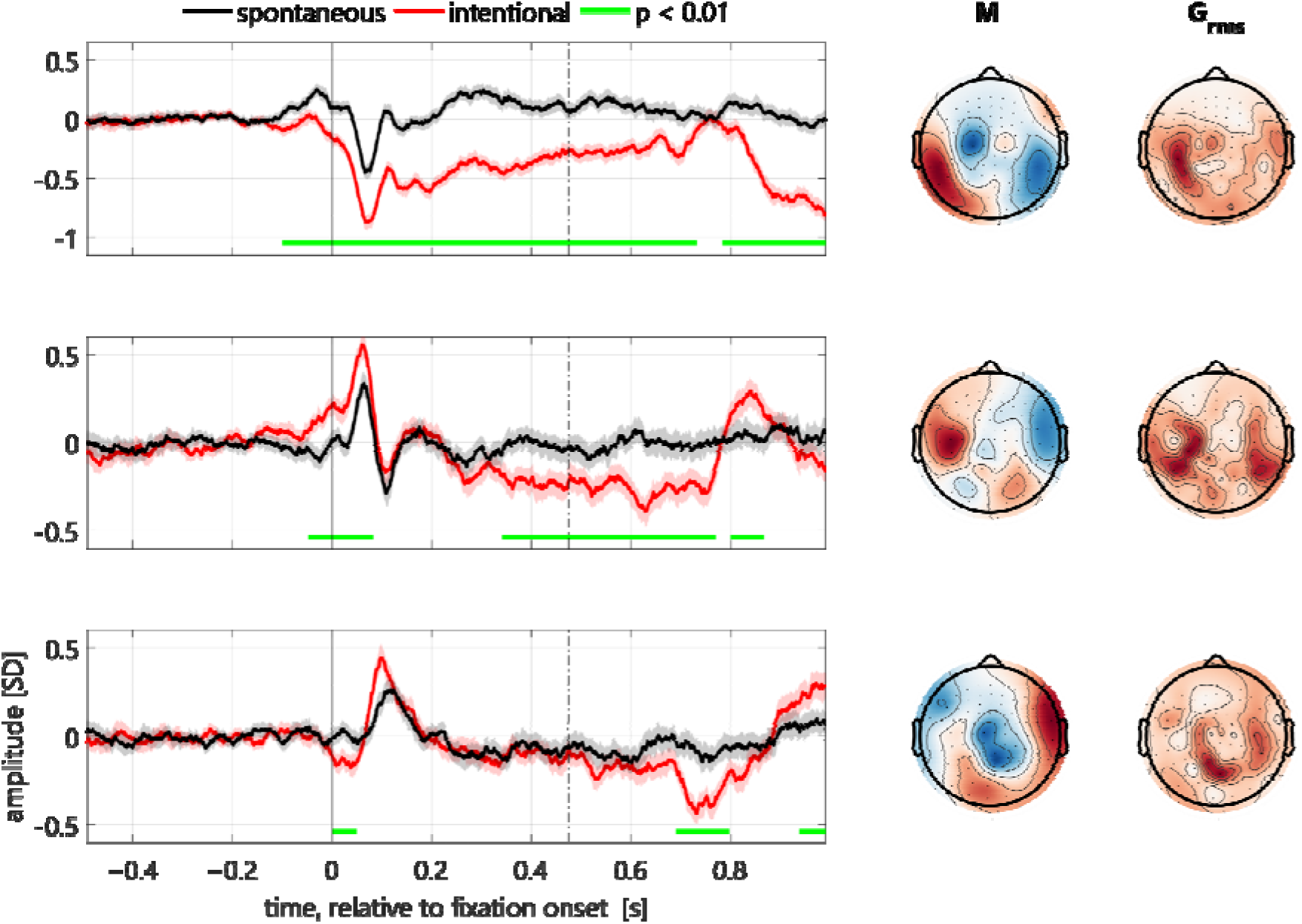
Event-related fields of the three significant spatial components. Left panel: median time-courses of components with 95% bootstrap (n = 1000) confidence interval of the median shown as colored shade. Right panel: sensor-level maps for the weights of forward models of the components projected onto magnetometers (M), and the root mean square of planar gradiometers (G_rms_). Vertical lines at 0 and 0.5 s show dwell onset and approximate feedback time, respectively.

The first and most significant component was primarily localized to the posterior region of the ventral surface of the temporal lobe bilaterally. In contrast, the second component exhibited a more distributed pattern across the ventral and dorsolateral cortical surfaces, with a notable cluster of sources in the orbitofrontal regions of both hemispheres (Figure 6, Table S3).

**Figure 6.**
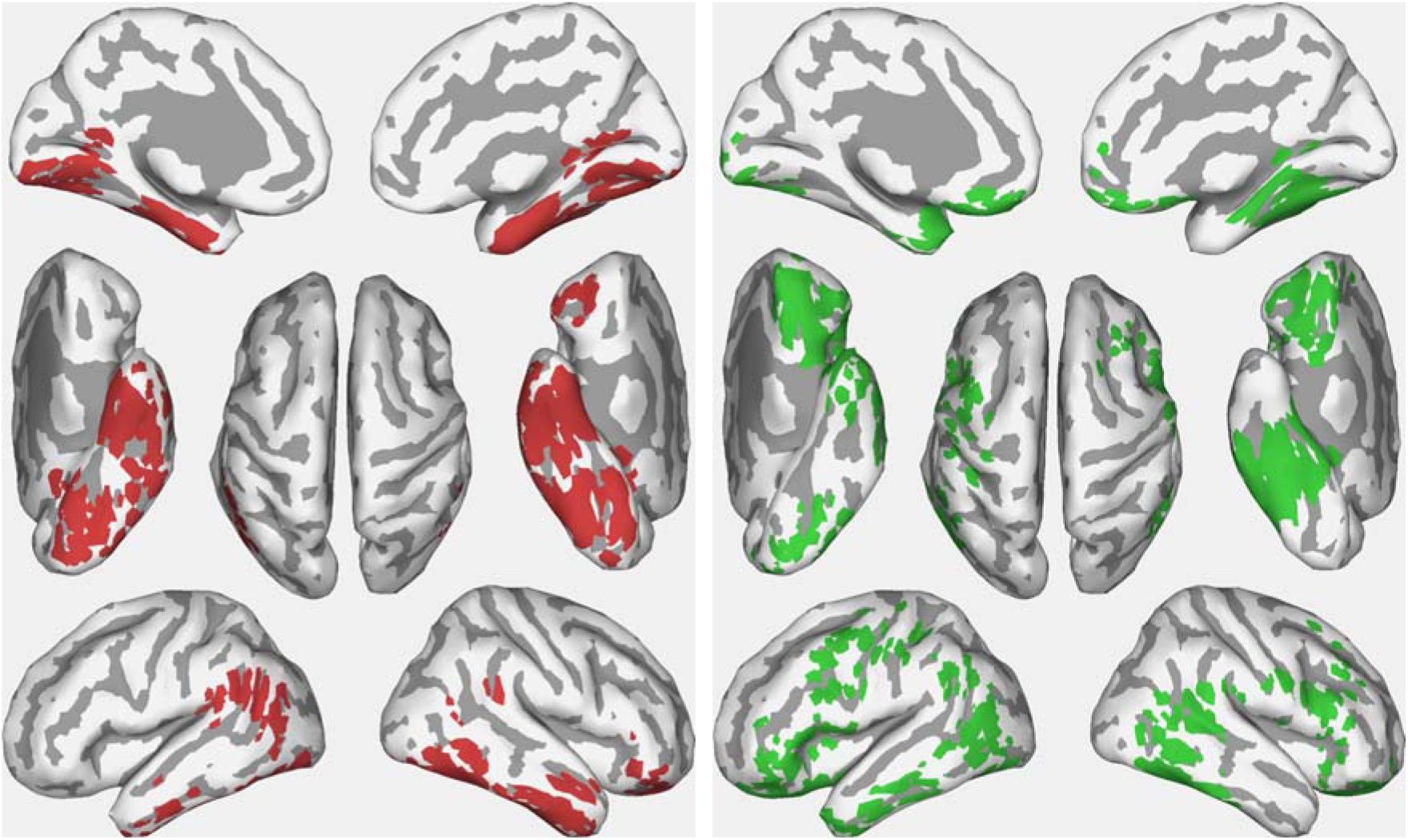
Grand average source representation of the two components of evoked activity that primarily distinguish intentional from spontaneous gaze dwells. All designations are as in Figure 4.

### 3.5. Control for eye movement artifact contribution

We checked if the ERF data were significantly affected by eye movement artifacts by averaging trials where dwell onset was preceded by a saccade with an amplitude greater than 2.6° (corresponding to a 200 px change in gaze coordinates) along the OX axis (labeled‘right’ and‘left’; Figure S3 and S4) and the OY axis (labeled‘up’ and‘down’; Figure S5 and S6). In intentional trials, these saccades accounted for 40.3% (OX) and 35.1% (OY) of all trials, while in spontaneous trials, they represented 26.6% (OX) and 24.7% (OY). The figures revealed no evidence of evoked component behavior that could be attributed to eye movement artifacts. Although large saccades did influence the time course of evoked components (see left panels on Figures S3-S6), they did not affect the difference between intentional and spontaneous responses (see right panels on the figures). Moreover, differences in oculomotor activity between spontaneous and intentional dwells (see Figure 2) could not account for the differences in MEG components, as they emerged after 300 ms—later than the development of MEG responses (see Figures 3 and 5).

## 4. Discussion

In this study, we, for the first time, utilized gaze-based human-computer interaction technology to examine the neural mechanisms that support the voluntary suppression of eye movements without external guidance.

Previous human neuroimaging studies on eye-movement suppression have employed external stop signals, either before target presentation (as in the antisaccade task (Munoz & Everling, 2004)) or after target onset (as in the countermanding paradigm (Kornylo et al., 2003)). Instead, our participants issued a gaze-based selection command by deliberately sustaining an extended fixation on a self-selected location on the game board. Our paradigm isolates the voluntary suppression of eye movements, removing additional processes related to stop-signal processing, redirection of attention, or preparation for a subsequent goal-directed saccade. By contrasting intentional and spontaneous fixations of extended duration, we were able to directly assess distinct characteristics of oculomotor activity, fixation-evoked responses, and oscillatory neural activity that specifically emerge during intentional gaze dwelling. Importantly, this approach reveals how intentional gaze withholding, in contrast to spontaneous dwelling, involves both top-down attentional control mechanisms and concurrent suppression within the oculomotor circuitry responsible for eye-movement initiation and planning, supporting a sustained focus on a defined spatial location.

In a broader perspective, the proposed contrast of intentionally vs. spontaneously prolonged gazes can be considered as a model for studying voluntary control and intention formation in general. Spontaneous eye movements and dwells are almost exclusively out of conscious control, unlike behaviors based on skeletal or even mimic muscle activity, and therefore may serve as a perfect control for voluntarily and consciously controlled intentional gaze behaviors in cognitive neuroimaging studies.

### 4.1. Oculomotor behavior characterizing intentional dwells

At the level of oculomotor behavior, we observed a notable difference between intentional and spontaneously occurring gaze dwells. Although both types showed strong suppression of eye movements during the first 200 ms after fixation onset, spontaneous dwells subsequently exhibited gradual gaze drift away from the fixation center (Figure 2B). In contrast, intentional gaze drift remained confined to a small (< 0.2°) spatial locus around the fixation center. Since smooth changes in eye position during natural fixations are typically involuntary (Krauzlis et al., 2017), this difference may suggest an active cortical control mechanism during intentional gaze dwelling, possibly enhancing precision for point-and-select actions via gaze. This inhibitory control appears to anchor the eyes firmly on the selected target, counteracting ocular drift, reducing discrepancies between eye and target position, and potentially preventing shifts in covert visual attention to peripheral locations, which are often accompanied by microsaccades and slower ocular drifts (Hafed & Clark, 2002). This mechanism likely supports the precision and stability necessary for intentional gaze-stopping actions. The coupling of oculomotor behavior with MEG recordings in our study offers some insights into the neural underpinnings of this control system.

Based on the observations made in this study, a separate line of research has been already started, where the *EyeLines* paradigm was used to extract intention-related features of oculomotor activity which differentiated intentional and spontaneous gaze dwells (Shevtsova et al., submitted; Shevtsova et al., 2023).

### 4.2. Neural concomitants of intentional dwells

Research on humans and other primates suggests that suppressing spontaneous oculomotor behaviors relies on proactive inhibition within the frontoparietal saccade network (Johnston et al., 2019). The specific properties of cortical oscillations associated with intentional versus spontaneous gaze dwellings in our study (Figure 3, 4) align well with the functional inhibition of frontal cortical centers responsible for eye movement initiation and planning.

Firstly, we observed increased alpha and beta power during intentional vs spontaneous dwellings (Figure 3, 4), which likely indicated functional inhibition in the underlying cortical circuitry. In human EEG and MEG studies, alpha power increases across different task contexts are commonly considered a marker of selective inhibition in cortical areas involved (e.g., Jensen & Mazaheri, 2010; Rihs et al., 2007), including those associated with eye movement planning and execution (Hwang et al., 2014). Supporting this, animal studies indicate that higher alpha power correlates with reduced neuronal firing—a hallmark of functional inhibition (Haegens et al., 2011).

Secondly, the topography of the differential alpha-beta band power increase was mainly bound to precentral scalp regions (Figure 3), suggesting a focus on motor rather than perceptual suppression. Source localization (Figure 4) confirmed that this oscillatory activity encompassed sensorimotor, premotor, and dorsolateral prefrontal cortex (DLPFC) regions, particularly including the frontal eye field (FEF) within the dorsal premotor cortex (at the junction of the superior precentral sulcus and superior frontal sulcus) and the supplementary eye field (SEF) along the border between medial and lateral surfaces of the frontal cortex (Figure 4, the first and the second components respectively). Research in nonhuman primates and human imaging studies suggests that SEF and FEF both contribute to ocular decision-making and the initiation of visually guided behaviors (Yang & Heinen, 2014). Therefore, the‘functional inhibition’ observed in these regions during intentional dwellings— evident in alpha and beta synchronization—may help establish a “status quo” state for the oculomotor system (see (Engel & Fries, 2010)), potentially suppressing any shifts in gaze position.

Additionally, the relatively widespread alpha-beta increase we found in the frontal cortex during intentional gaze dwellings may arise from functional inhibition within a broader network of premotor and prefrontal cortical areas that contributes to the quieting of head and body movement, helping to keep the gaze within the designated spatial locus by minimizing disruptive head movements (Corneil et al., 2008). Notably, we did not observe significant intention-related modulations in the posterior cortical areas, specifically in the posterior cortical eye fields situated near the inferior parietal sulcus (IPS) that are involved in oculomotor behavior (Andersen, 1989). This suggests that the oculomotor processes supported by the IPS may have been equivalent between spontaneous and intentional gaze dwells.

Thirdly, the timing of induced alpha-beta synchronization during intentional gaze dwellings implied that this oscillatory activity began at least a couple of hundred milliseconds before gaze stabilization within the intended spatial locus (Figure 3, Figure S2), suggesting a proactive inhibition control of the cortical saccade initiation network.

The latter finding is generally consistent with previous literature, though prior neuroimaging research has primarily focused on the preparatory period of the tasks requiring suppression of the prepotent oculomotor response upon the external cue (see, e.g., (Brown et al., 2006)). In the anti-saccade (AS) task, participants are instructed either to direct their gaze toward an upcoming visual target or to suppress this reflexive response and instead shift their gaze voluntarily to the opposite location, depending on the cue provided. Although human EEG/MEG research on the preparatory phase of the AS task remains limited, recent MEG findings indicate that successful task performance is associated with increased preparatory alpha-band power in the frontal eye fields (FEF) following the cue and prior to target onset (Hwang et al., 2014, 2016). Remarkably, a similar task-dependent increase in alpha oscillations was observed in monkey FEF local field potentials during a preparatory period when the monkeys were cued to inhibit a prepotent oculomotor response (Johnston et al., 2019). In this task, the monkeys had to either direct their gaze toward a large, high-luminance stimulus or inhibit this response in favor of a saccade toward a smaller, low-luminance stimulus. When instructed to suppress a reflexive gaze shift, prominent pre-stimulus alpha-band activity appeared in deeper FEF layers, correlating with reduced neuronal firing in upper layers, suggesting that alpha-band rhythms in deep layers support proactive saccade inhibition.

Our results confirm and extend the previous literature, by showing that alpha-beta activity in the FEF and SEF, reflecting proactive, top-down inhibitory control, can arise without an external cue, solely driven by the intention to sustain gaze dwelling.

Alongside the induced oscillations, we observed distinctive evoked activity associated with intentional gaze withholding versus spontaneous gaze dwelling. The primary difference between these conditions in the fixation-onset-locked magnetic fields was not in transient ERF peaks but rather in a gradual DC shift in the evoked response waveform (Figure 5), suggesting sustained cortical activation associated with intentional gaze withholding. Literature suggests that such long-lasting evoked activation—extending from hundreds of milliseconds to several seconds in EEG/MEG—often emerges gradually in anticipation of predictable or relevant events and may be linked to attentional processing ((Pulvermüller & Grisoni, 2020); for review see (León-Cabrera et al., 2024)). In our study, intentional fixations were indeed associated with the anticipation of the feedback from the gaze-sensitive interface, which allowed participants to terminate gaze holding and shift their gaze to another location (see Methods). However, the timing of the slow deflection, which peaked approximately 200-300 ms prior to, rather than at, the anticipated event, suggests it does not reflect preparatory cortical activity (Kononowicz & Penney, 2016).

An alternative interpretation of the sustained neural activation evoked by intentional fixation is that it is attention-related. To use their gaze as a selection gesture, our participants needed not only to inhibit oculomotor behavior but also to concentrate and continuously focus on a self-selected spatial location on the game board. This steady focus was essential to effectively transmit a command to the computer, because it ensured that the gaze was kept in the “gaze-sensitive” area.

We propose that the sustained neural activation observed during intentional gaze dwelling reflects this voluntary spatial attention focusing on a specific spot. Based on its scalp distribution (Figure 5) and crude source localization (Figure 6), the cortical sources of this activity are primarily located in the inferior convexity of the cortex, specifically in the posterior inferior temporal cortex (ITC) and inferior frontal regions (see also Table S3 for components 1 and 2).

Recent neurophysiological studies provide evidence that persistent neuronal firing in the posterior ITC is strongly modulated by spatial attention. The researchers emphasized that the posterior ITC may function as a distinct system for spatial attention focus, directing it according to an individual’s internal objectives, thereby supporting cognitive programs relevant to goal-directed actions (Ramezanpour & Fallah, 2022; Sani et al., 2021; Stemmann & Freiwald, 2019). This system is thought to encode a “priority map” that determines which locations in the visual field are attended (Stemmann & Freiwald, 2019). Intentional fixations in our study may thus recruit this spatial attention system to sustain gaze on a specific point, reducing diversions from that location.

The sustained activation of the orbitofrontal cortex (OFC)/ventral prefrontal cortex (PFC) (Figure 6) aligns with the proposed role of goal-directed spatial attention in intentional gaze fixations. In both primate neurophysiology and human neuroimaging studies, this region is recognized as part of the neural circuitry involved in oculomotor control, particularly in maintaining goal-relevant information (Fielding et al., 2015; Xu et al., 2017). During intentional gaze fixation, sustained OFC/ventral PFC activity may facilitate the registration of the intention to inhibit gaze shifts, potentially interacting with the ventral temporal attentional system to exert inhibitory control over eye movements.

Overall, our neurophysiological results suggest distinct functional roles for sustained evoked activation and induced oscillations during intentional gaze fixations, which were used to trigger a selection operation in the context of interaction with a computer. The prolonged evoked activation in the posterior inferior temporal cortex (ITC) and ventral prefrontal cortex (PFC), time-locked to fixation onset, appears to reflect the robust control of spatial attention on a specific location, thereby supporting the suppression of oculomotor behavior. Meanwhile, alpha-beta band oscillatory activity in the frontal eye fields reflects the strength and efficacy of this inhibitory control.

### 4.3. Limitations

Our study has several limitations. First, the precise timing of intention formation to select a specific spatial location on the game board could not be determined. While our results suggest that intention formation may be extended, encompassing both pre-and post-onset intervals, its timing likely varies across trials, potentially smearing the observed temporal patterns. Second, in our experiment design intentional dwells were different from the spontaneous ones not only by a different degree of intentionality but also by different eye movement characteristics in the latter portion of the dwell and especially immediately after it, and by the presence of planned action to make a saccade to the confirmation location. It seems highly unlikely that these differences could affect our results, but they may restrict possible areas of the application of the proposed methodology. However, at least the issue of the saccade planning in case of intentional dwells can be solved by using more advanced gaze-based control techniques not requiring confirmation or similar actions, such as gaze-based control enhanced by machine learning and context-based predictions (Shevtsova et al., submitted; Shevtsova et al., 2023). Third, the absence of individual MRI data required us to use a standardized brain template, which may have introduced inaccuracies in localization. Fourth, performing the GED analysis at the group level rather than individually might have similarly affected the precision of our results. Despite these limitations, we believe they do not substantially impact the main findings of this study.

### 4.4. Intentionally prolonged gaze in natural behavior

The presented methodology of contrasting intentional and spontaneous gaze behavior observed in the context of using gaze-based interaction technology can be considered as being closer to the natural use of gaze than the experimental paradigms requiring significant efforts to modify reflective behavior.

More specifically, the origin of the ability to use gaze voluntarily may be related to its use in social interaction, where it is typically used without significant effort. Gaze-based social interaction is increasingly studied in behavior patterns like direct gaze (Adams & Kleck, 2005; Hamilton, 2016; Senju & Hasegawa, 2005) and joint attention (Chevalier et al., 2020; Siposova & Carpenter, 2019; Tomasello & Farrar, 1986). However, the main focus of such studies is often on gaze perception, and research on the active, “sending” side of gaze-based social interaction has only recently begun (Fedorova et al., 2015; Gobel et al., 2015; Hessels et al., 2019; Jarick & Bencic, 2019; Riechelmann et al., 2021; Risko et al., 2016). In a more effortful task of avoiding one‘s partner’s gaze the cortical activation pattern was similar to that observed in antisaccade tasks (Cavallo et al., 2015). To our knowledge, brain activity has not been yet studied in such social paradigms where gaze is under low-effort voluntary control. Such studies could likely bring new insights into voluntary gaze control, but gaze-based interaction with technical devices has some advantages as an experimental paradigm, as providing more strict control over the factors involved in an experiment.

### 4.5. Prospective application in hybrid eye-brain-computer interfaces (EBCIs)

Estimating the intentionality of gaze dwells using EEG/MEG could improve gaze-based interaction by preventing unintended computer responses to similarly prolonged spontaneous fixations, thereby enabling a more seamless user-computer interaction (Ihme & Zander, 2011; Protzak et al., 2013; Shishkin et al., 2016b). Shishkin et al. (2016b) proposed the term “eye-brain-computer interface” (EBCI) to distinguish this hybrid system from mechanistically combined BCIs and gaze interfaces. EEG studies have identified the expectancy wave (E-wave, also referred to as stimulus-preceding negativity, SPN, or non-motor contingent negative variation, CNV) as a marker of intentionality, based on its presence in intentional but not spontaneous gaze dwells, in different paradigms: in the EyeLines game (Shishkin et al., 2016b), in a task involving a selection of moving targets with smooth pursuit eye movement (“dynamical dwells”) (Zhao et al., 2021), in target selection in extended reality (XR) (Reddy et al., 2024). Nuzhdin et al. (2017) demonstrated that a passive BCI could use this marker to classify gaze dwells online, though the system’s performance fell short of practical application. A key limitation is that the E-wave is not specific to intentional gazes, as it can reflect the expectation of interface feedback even in the absence of intent. In one study (Shishkin et al., 2016a) classification of intentional and spontaneous dwells was improved using features of oscillatory EEG components, which could be, in principle, more specific to intentional gaze dwells. These results, however, could be biased because of possible contamination in the early part of the studied EEG interval, where a large saccade often occurred in intentional dwells due to specific organization of the experimental design.

Notably, in the current study, none of the sustained phase-locked activity components differentiating intentional from spontaneous gaze dwells exhibited the characteristic temporal profile of the E-wave observed in prior EEG studies. While this discrepancy may result from differences in how MEG and EEG capture neural activity, it also suggests that the intentionality-related components identified here may reflect the deliberate focusing of gaze on a specific spatial location, rather than merely anticipation of feedback.

An important advantage of the present study over previous ones is the use of a modified paradigm, where labeling of intentional and spontaneous dwells was based on gazing at the confirmation screen button after the intentional selection, not before it. In this paradigm, eye movements near the onset of intentional and spontaneous dwells are similar. Furthermore, the use of MEG instead of EEG enabled improved spatial source separation. These findings can guide future research employing deep neural networks to classify spontaneous and intentional dwells based on MEG data (Ovchinnikova et al., 2021), providing a stronger foundation for more reliable classification—an essential step toward the practical application of hybrid Eye-Brain-Computer Interfaces.

While MEG is currently too costly for widespread use in commercial BCIs, recent advancements in optically pumped magnetometers (OPMs) (Boto et al., 2018; Boto et al., 2017) and other compact, room-temperature MEG sensors (Koshev et al., 2021; Petrenko et al., 2021) have opened a perspective for MEG-based BCI applications (Iivanainen et al., 2023; Ji et al., 2024). These technological achievements could enable the integration of MEG-informed intentional gaze detection into practical assistive technologies.

Furthermore, it may be valuable to explore EEG components corresponding to the MEG components identified in this study, with the goal of developing similar applications using EEG systems, which are more portable and cost-effective.

### 4.6. Implications for control conditions in neuroimaging

Maintaining gaze on a fixation cross or similar object is a common “baseline” condition in neuroimaging studies. Although our design required shorter fixation times, even this modest demand influenced brain activity, suggesting that longer fixations might further impact neural processes. These findings add to concerns (e.g., (Northoff et al., 2010)) that fixation cross conditions may not be entirely neutral and could influence brain activity. Further research is needed to explore this issue.

## 5. Conclusions

This study employed a novel experimental paradigm combining gaze-based human-computer interaction with 306-channel magnetoencephalography (MEG) to explore the neural mechanisms of voluntary gaze control. Participants played a video game using gaze-based actions, allowing us to compare intentional and spontaneous gaze dwells in a naturalistic context. Our analysis revealed alpha-beta band synchronization (8–30 Hz) within the frontal cortex, beginning prior to an intentional dwell’s onset and peaking shortly afterward, likely reflecting proactive inhibitory mechanisms for saccadic suppression. Additionally, sustained activation in the inferior temporal and orbitofrontal cortices persisted throughout the duration of intentional gaze dwells, possibly supporting focused spatial attention during intentional fixation. These findings provide new insights into the cortical dynamics of voluntary gaze control, with implications for improving gaze-based interaction technologies. Future research could refine cortical activation mapping to enhance understanding and applications of voluntary gaze mechanisms.

## Acknowledgments

Authors thank A.O. Prokofyev and A.S. Yashin for their help in running experiments and MEG data preprocessing.

This work was supported by the Russian Science Foundation (grant number 22-19-00528).

The study was conducted at the unique research facility “Center for Neurocognitive Research (MEG-Center)” of MSUPE.

## Declaration of generative AI and AI-assisted technologies in the writing process

During the preparation of this work the authors used ChatGPT (OpenAI) in order to improve the readability and language of the manuscript. After using this tool, the authors reviewed and edited the content as needed and take full responsibility for the content of the published article.

## Footnotes

1 We use the term *dwell* for the events of relatively long stay of gaze in a designated area (area of interest). Such events are often called *fixations*, especially in the literature on gaze-based interaction, but they do not need to fit strict definitions of an eye fixation used in the eye movement literature. Moreover, a dwell may include more than one fixation (in strict definition) and other kinds of eye movement.

