## Supplementary materials for "Intentionally vs. Spontaneously Prolonged Gaze: A MEG Study of Active Gaze-Based Interaction" (from another version of this preprint: version 1, 2024-12-16)

### Supplementary Material

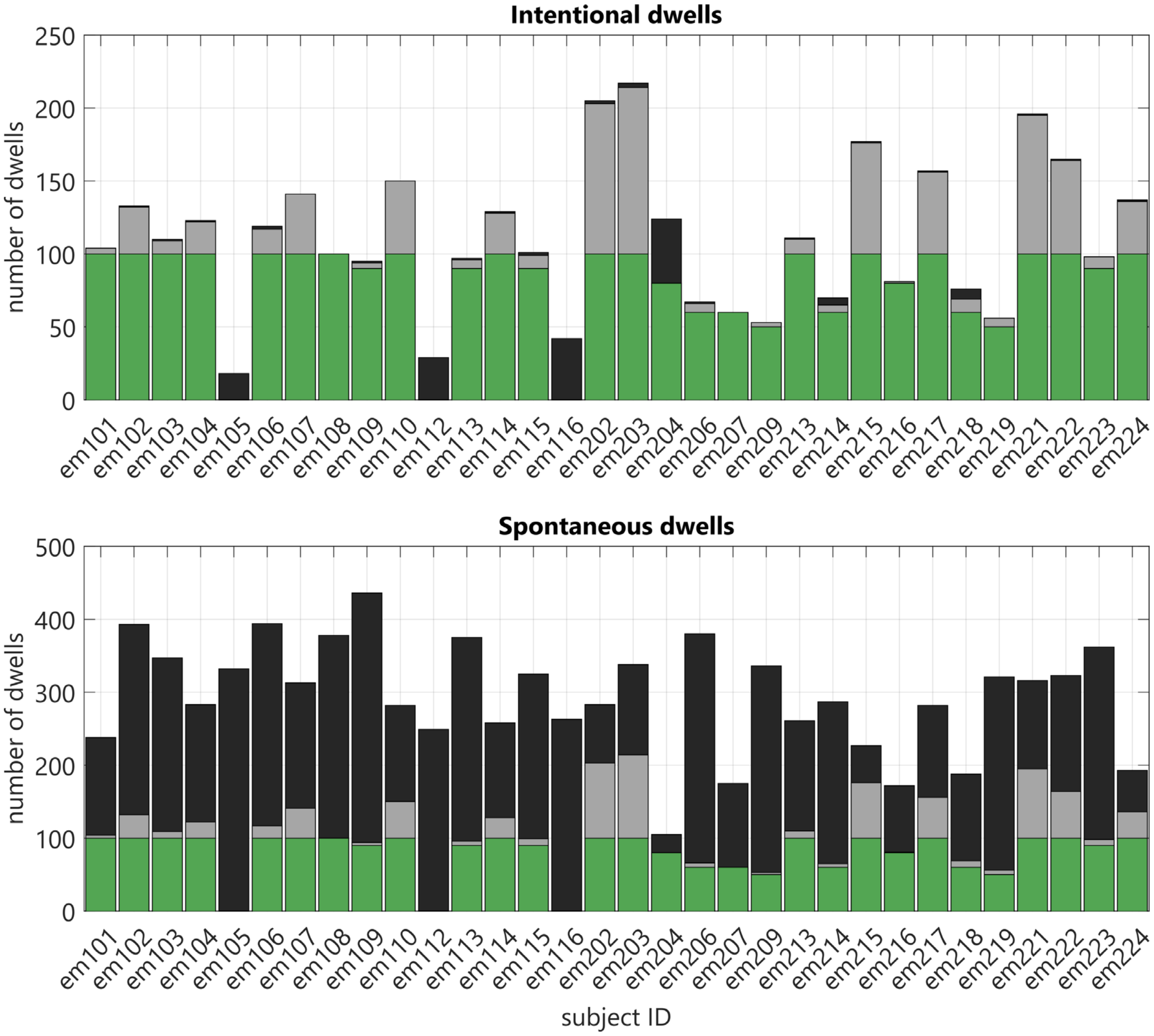

Figure S1. Progression from the raw set of dwell-related data to the finally used dataset. Total bar length corresponds to the number of dwells per class (intentional, spontaneous) and participant; black – initial dataset; grey – removed to equalize classes or because of low MEG data quality; green – removed to equalize across participants.

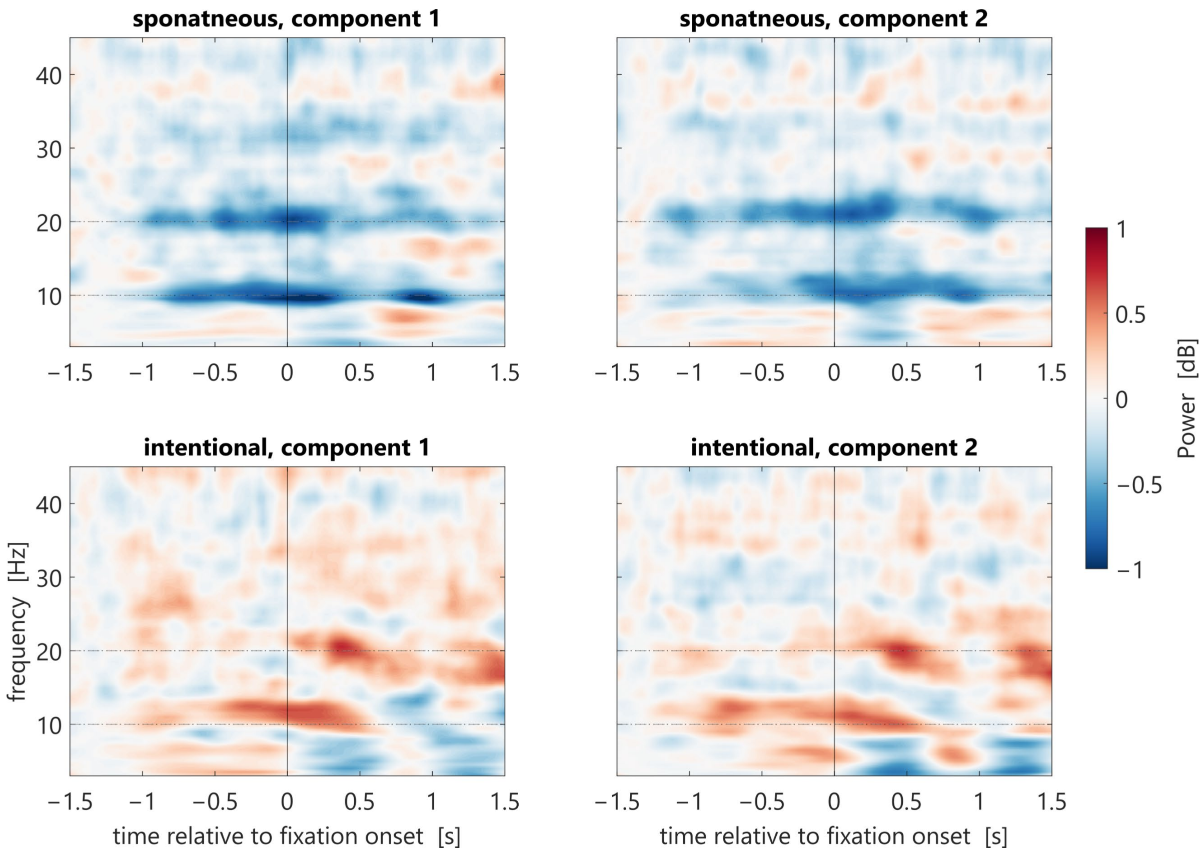

Figure S2. Time-frequency maps reflecting baseline normalized power changes for each component, plotted for spontaneous and intentional dwells separately. The baseline interval corresponds to [-1.5 – -1.2] s relative to fixation onset.

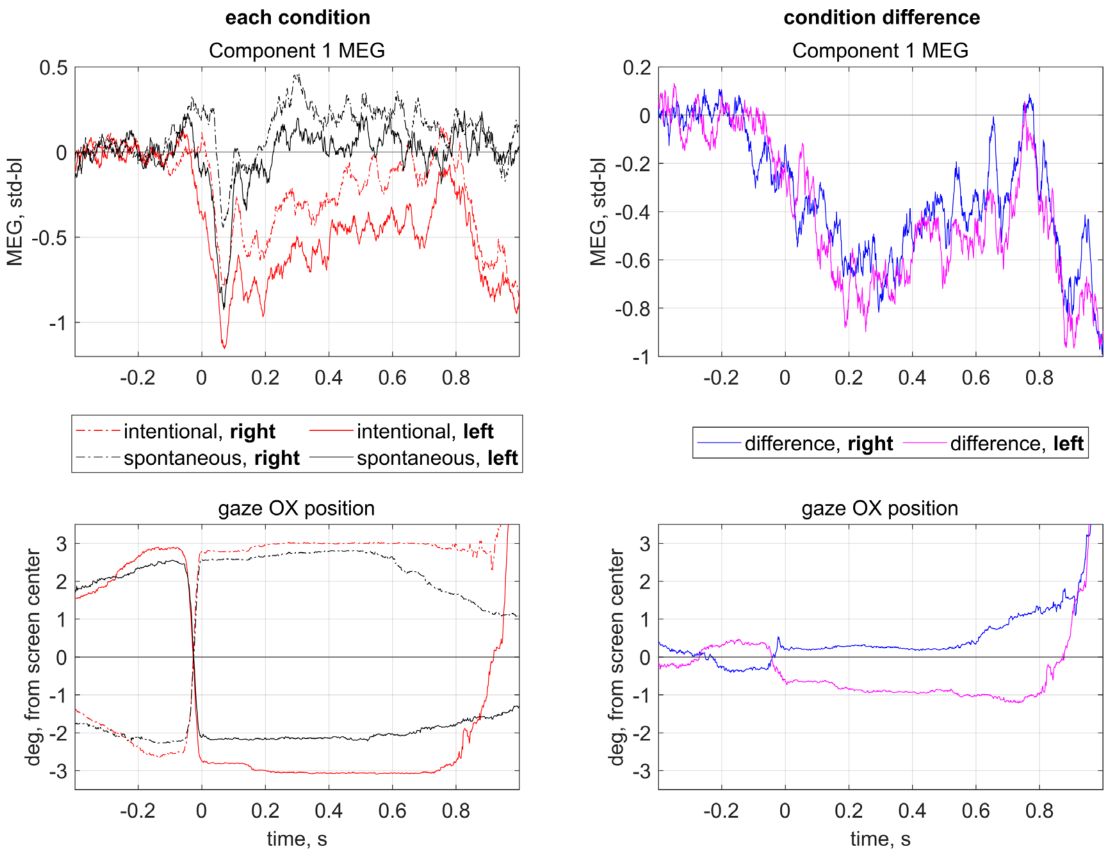

Figure S3. Left panel – (Top) Event-related fields (ERF) for the first significant spatial component averaged across all epochs preceded by saccades larger than 2.6° along the OX Solid lines represent leftward saccades, while dotted lines represent rightward saccades. Values are presented as standard deviations from the baseline (-300 to -100 ms). (Bottom) the same data for the relative gaze coordinates. Right panel – The difference in ERF (top) and gaze coordinates (bottom) between "spontaneous" and "intentional" dwell conditions.

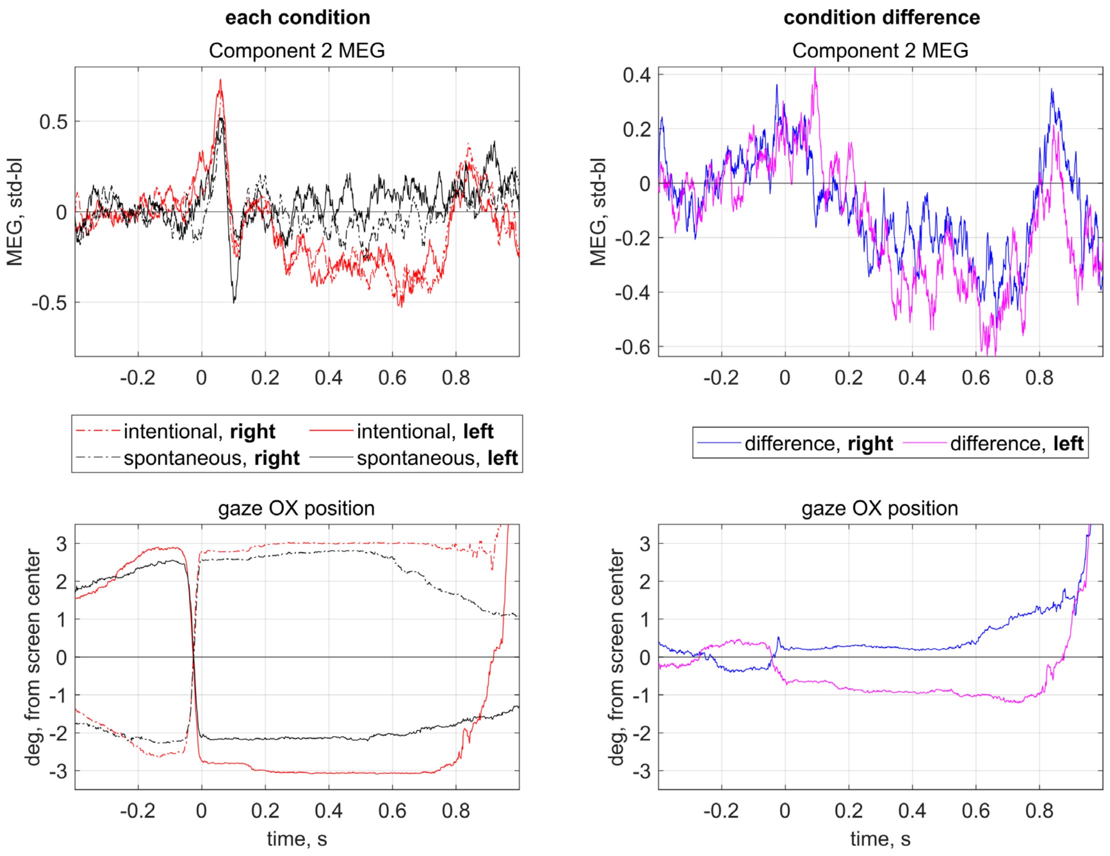

Figure S4. Same as in Figure S3 but for the second significant spatial component.

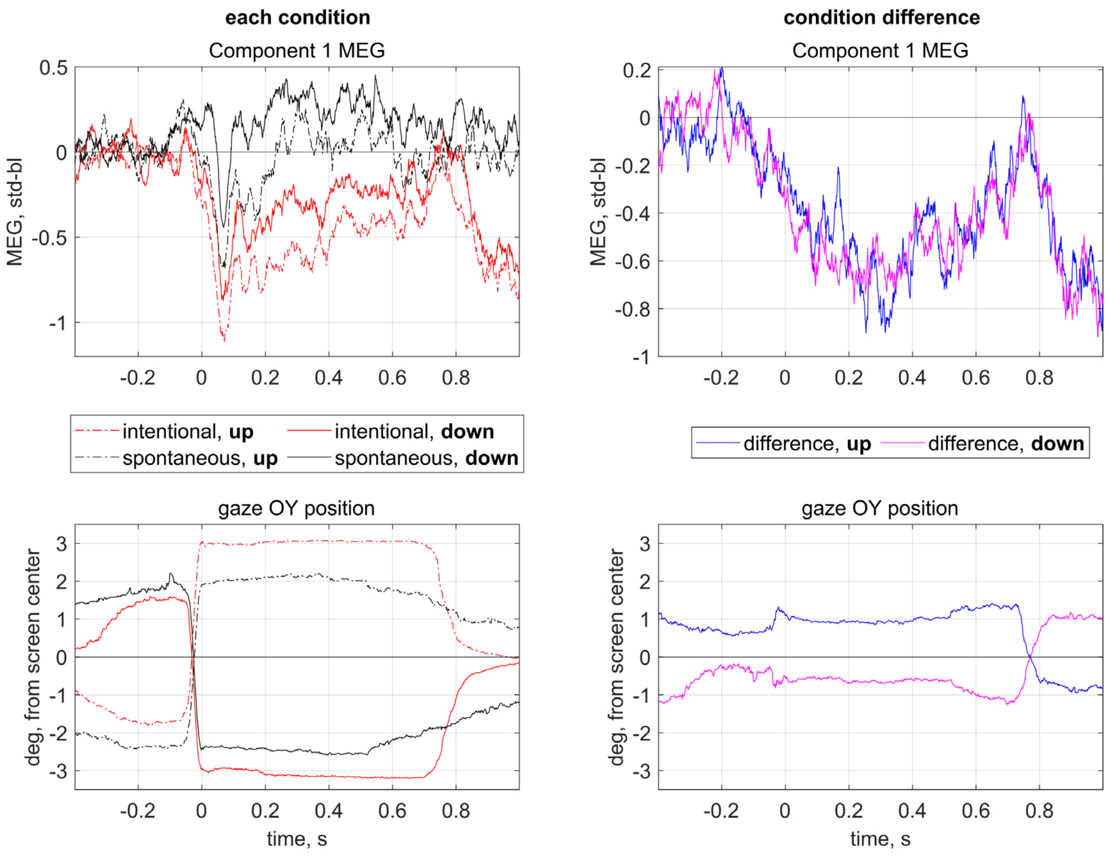

Figure S5. Left panel – (Top) Event-Related Fields (ERFs) for two significant spatial components, averaged across all epochs preceded by saccades greater than 2.6° along the OY axis. Solid lines represent leftward saccades, while dotted lines represent rightward saccades. Values are presented as standard deviations from the baseline (-300 to -100 ms). (Bottom) The same data is illustrated for relative gaze coordinates. Right panel – Displays the difference in ERF (top) and gaze coordinates (bottom) between the "spontaneous" and "intentional" dwell conditions.

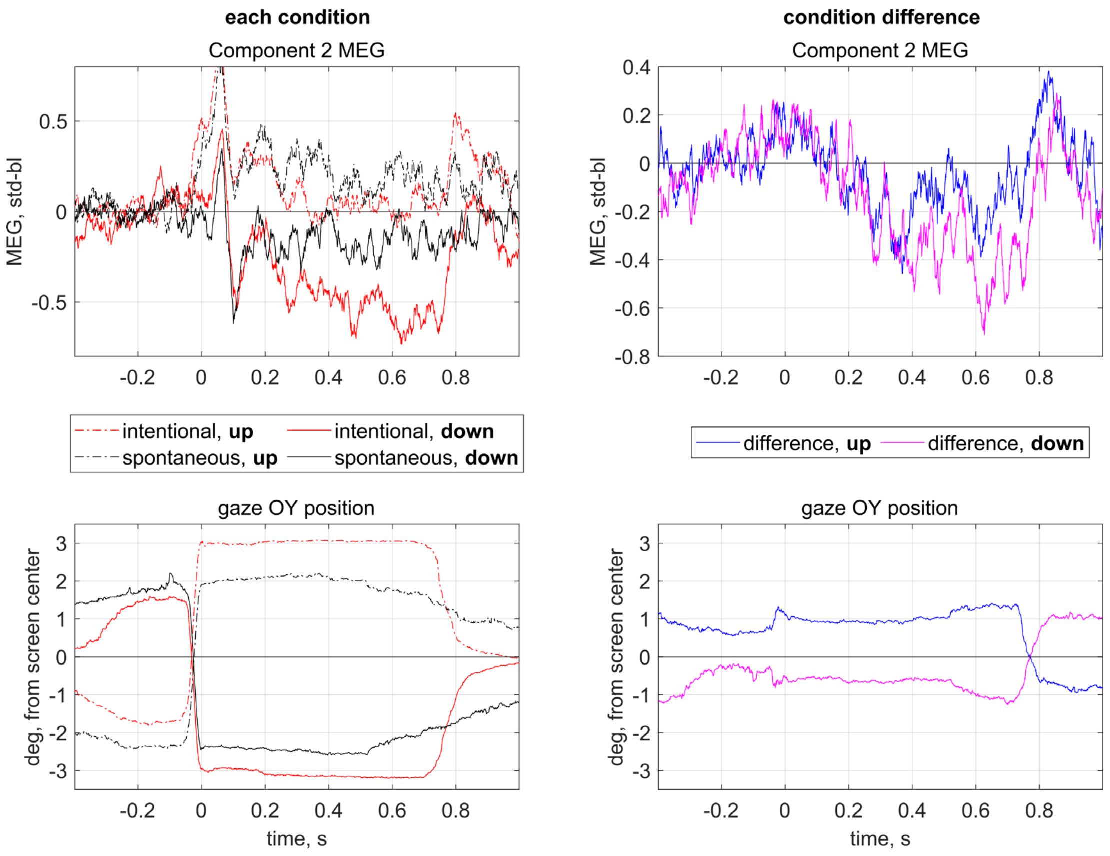

Figure S6. Same as in Figure S5 but for the second significant spatial component.

Table S1. Number of selections and dwells per participants.

| subj ID | per game mean | | | all games total | | | time between moves, s | | |
|---|---|---|---|---|---|---|---|---|---|
| | all selections | unconfirm. selections | confirm. selections | all selections | unconfirm. selections | confirm. selections | all selections | unconfirm. selections | confirm. selections |
| em101 | 88.1 | 47.8 | 40.4 | 705 | 382 | 323 | 3.44 | 7.28 | 5.97 |
| em102 | 95.6 | 61.3 | 34.4 | 765 | 490 | 275 | 3.13 | 8.29 | 4.78 |
| em103 | 101.4 | 61.9 | 39.5 | 811 | 495 | 316 | 2.99 | 7.58 | 4.83 |
| em104 | 78.9 | 45.3 | 33.6 | 631 | 362 | 269 | 3.78 | 8.35 | 6.39 |
| em105 | 96.9 | 65.4 | 31.5 | 775 | 523 | 252 | 3.13 | 9.63 | 4.52 |
| em106 | 113.0 | 74.4 | 38.6 | 904 | 595 | 309 | 2.68 | 7.76 | 4.02 |
| em107 | 97.4 | 54.1 | 43.3 | 779 | 433 | 346 | 3.13 | 6.83 | 5.52 |
| em108 | 101.6 | 65.4 | 36.3 | 813 | 523 | 290 | 2.93 | 7.98 | 4.45 |
| em109 | 108.5 | 76.6 | 31.9 | 868 | 613 | 255 | 2.63 | 8.78 | 3.63 |
| em110 | 90.0 | 47.8 | 42.3 | 720 | 382 | 338 | 3.28 | 6.85 | 6.03 |
| em111 | 94.0 | 65.4 | 28.6 | 752 | 523 | 229 | 3.22 | 10.42 | 4.57 |
| em112 | 66.1 | 40.6 | 25.5 | 529 | 325 | 204 | 4.61 | 11.2 | 7.37 |
| em113 | 100.6 | 67.5 | 33.1 | 805 | 540 | 265 | 2.97 | 8.90 | 4.34 |
| em117 | 89.4 | 44.6 | 44.8 | 715 | 357 | 358 | 3.28 | 6.37 | 6.36 |
| em115 | 95.8 | 56.9 | 38.9 | 766 | 455 | 311 | 3.15 | 7.63 | 5.20 |
| em116 | 77.5 | 57.0 | 20.5 | 620 | 456 | 164 | 3.58 | 13.04 | 4.77 |
| em202 | 83.1 | 45.6 | 37.5 | 665 | 365 | 300 | 3.59 | 7.58 | 6.36 |
| em203 | 102.8 | 53.3 | 49.5 | 822 | 426 | 396 | 2.97 | 6.06 | 5.66 |
| em204 | 72.5 | 26.6 | 45.9 | 580 | 213 | 367 | 4.18 | 6.49 | 11.42 |
| em206 | 97.0 | 70.6 | 26.4 | 776 | 565 | 211 | 2.76 | 9.48 | 3.69 |
| em207 | 65.5 | 36.5 | 29.0 | 524 | 292 | 232 | 4.61 | 9.35 | 8.21 |
| em209 | 90.6 | 61.3 | 29.2 | 815 | 552 | 263 | 3.24 | 9.57 | 4.74 |
| em213 | 86.9 | 48.4 | 38.5 | 695 | 387 | 308 | 3.40 | 7.64 | 5.77 |
| em214 | 74.2 | 44.7 | 29.6 | 668 | 402 | 266 | 3.32 | 8.21 | 5.29 |
| em215 | 90.1 | 41.5 | 48.6 | 721 | 332 | 389 | 3.11 | 5.69 | 6.43 |
| em216 | 72.3 | 39.4 | 32.9 | 506 | 276 | 230 | 4.16 | 8.77 | 7.42 |
| em217 | 90.6 | 50.9 | 39.8 | 725 | 407 | 318 | 3.34 | 7.30 | 5.93 |
| em218 | 77.1 | 44.3 | 32.9 | 617 | 354 | 263 | 3.87 | 8.90 | 6.55 |
| em219 | 89.1 | 59.4 | 29.8 | 713 | 475 | 238 | 3.36 | 9.93 | 4.93 |
| em221 | 106.9 | 56.5 | 50.4 | 855 | 452 | 403 | 2.82 | 5.92 | 5.01 |
| em222 | 90.0 | 49.3 | 40.7 | 810 | 444 | 366 | 2.96 | 6.40 | 5.15 |
| em223 | 81.4 | 57.4 | 24.0 | 651 | 459 | 192 | 3.71 | 12.37 | 5.24 |
| em224 | 88.9 | 46.3 | 42.6 | 711 | 370 | 341 | 3.35 | 6.52 | 6.18 |
| Mean±SD | 89.5±11.9 | 53.5±11.5 | 36.1±7.6 | 722±100 | 431±95 | 291±61 | 3.35±0.50 | 8.27±1.78 | 5.66±1.48 |

Table S2. Anatomic areas explaining the first and second induced field components.

| | Centroid [x, y, z] mm, MNI coordinates | Number of vertices | Anatomical location of centroid, closest Brodmann area |
|---|---|---|---|
| **Induced field component 1** | | | |
| Dorsal side (frontal, central, parietal) | L [-40 -13 50]<br>R [47 -15 45] | L 262<br>R 239 | L premotor, supplementary motor (BA6)<br>R primary motor (BA4) |
| **Induced field component 2** | | | |
| Dorsal side (frontal, central, parietal) | L [-38 -15 52]<br>R [41 -18 49] | L 249<br>R 254 | L premotor, supplementary motor (BA6)<br>R primary motor (BA4) |

Table S3. Anatomic areas explaining the first and second evoked field components.

| | Centroid [x, y, z] mm, MNI coordinates | Number of vertices | Anatomical location of centroid, closest Brodmann area |
|---|---|---|---|
| **Evoked field component 1** | | | |
| Ventro-temporal | L [-29 -51 -20]<br>R [36 -40 -23] | L 190<br>R 281 | L fusiform (BA37)<br>R fusiform (BA37) |
| Tempo-parietal | L [-52 -55 18]<br>R [57 -23 -14] | L 109<br>R 57 | L angular gyrus (BA39)<br>R middle temporal gyrus (BA21) |
| **Evoked field component 2** | | | |
| Ventro-temporal | L [-37 -46 -23]<br>R [39 -50 -19] | L 90<br>R 171 | L fusiform (BA37)<br>R fusiform (BA37) |
| Tempo-parietal | L [-47 -31 -5]<br>R [55 -55 7] | L 137<br>R 77 | L middle temporal gyrus (BA21)<br>R fusiform (BA37) |
| Fronto-central | L [-45 -5 42]<br>R [50 8 29] | L 76<br>R 44 | L premotor, supplementary motor (BA6)<br>R premotor, supplementary motor (BA6) |
| Orbito-frontal | L [-15 33 -21]<br>R [15 38 -20] | L 98<br>R 81 | L orbitofrontal (BA11)<br>R orbitofrontal (BA11) |
| Opercular | L [-47 -25 1]<br>R [50 19 15] | L 84<br>R 54 | L primary auditory (BA41)<br>R Broca (BA44) |
